# Characterization of Cytochrome P450-Mediated Metabolism of Δ^8^-tetrahydrocannabinol *in vitro*

**DOI:** 10.64898/2026.10.12.758575

**Authors:** Keiann T. Simon, Yue Winnie Wen, Yuqi Dong, Aurora K. Authement, Jiayao Chen, Kenneth E. Thummel, Nina Isoherranen

**Author notes:** **Contact Information for Corresponding Author:**Nina Isoherranen, 1959 Pacific Ave, HSB H-272N, Seattle, WA 98195 (206)543-0829.

## Abstract

Δ^8^-tetrahydrocannabinol (Δ^8^-THC) is a pharmacologically active isomer of Δ^9^-THC, the main psychoactive component of cannabis. Δ^8^-THC use has increased rapidly, but the metabolism of Δ^8^-THC has been incompletely characterized. This study identified metabolites of Δ^8^-THC formed by cytochrome P450 (CYP) enzymes in human liver microsomes (HLM), assigned the main CYPs responsible for the metabolic pathways, and evaluated the impact of genetic polymorphisms on Δ^8^-THC metabolism *in vitro*. Five monohydroxylated Δ^8^-THC metabolites, identified as 11-OH-Δ^8^-THC, two aliphatic side-chain hydroxylated products (M1, M2), and two ring hydroxylated products (M3, M4) were detected in HLM. 11-OH-Δ^8^-THC was the most abundant metabolite. Unbound binding constants for metabolite formation in HLM ranged from 3.6 nM to >121 nM, suggesting multiple enzymes contribute to Δ^8^-THC metabolism. The contributing enzymes were identified using recombinant CYPs, inhibition studies, and correlation analyses. 11-OH-Δ^8^-THC was predominantly formed by CYP2C9 with a minor contribution from CYP2C19. CYP2C9 genotype-based activity scores across donors correlated with 11-OH-Δ^8^-THC formation (p = 0.0315). An 80% decrease in 11-OH-Δ^8^-THC formation clearance was predicted in CYP2C9*3/*3 carriers. The ring-hydroxylated M3 and M4 were mainly formed by CYP3A. CYP2D6, CYP1A2, and CYP3A4 activities were associated with formation of the pentyl side-chain oxidized metabolites M1 and M2. Although C11 hydroxylation appeared more important metabolic pathway for Δ^8^-THC than for Δ^9^-THC, the catalytic efficiency of CYP2C9 towards Δ^8^-THC was lower than towards Δ^9^-THC. In the presence of CYP2C9 inhibitors or genetic polymorphisms, Δ^8^-THC clearance is likely mediated by CYP3A4 and CYP2C19.

**Significance statement:** This study shows that 11-OH-Δ^8^-THC, a psychoactive metabolite, is the primary oxidation product of Δ^8^-THC in HLM and that CYP2C9 predominantly catalyzes its formation. This work expands the limited literature on Δ^8^-THC metabolism, and shows that reduced-function CYP2C9 variants substantially impair formation of 11-OH-Δ^8^-THC. The characterization of Δ^8^-THC metabolism in HLM provides the foundation for risk assessment of drug-drug interactions and establishes the metabolic profile of Δ^8^-THC in the human liver.

## 1. Introduction

Δ^8^-tetrahydrocannabinol (Δ^8^-THC) is a minor constituent in *Cannabis sativa*, typically present only at trace levels.^1^ Δ^8^-THC is mainly obtained through chemical isomerization of cannabidiol extracted from hemp. Δ^8^-THC is structurally nearly identical to the better characterized Δ^9^-THC, the main psychoactive component of cannabis (**Figure 1**), but users report a preference for Δ^8^-THC over Δ^9^-THC.^2,3^ A substantial increase in Δ^8^-THC exposures has been reported by poison control centers, raising concerns about the safety of widespread Δ^8^-THC use.^4^ Despite the increasing use and potential risks of Δ^8^-THC, the metabolism and disposition of Δ^8^-THC has been incompletely characterized.

**Figure 1:**
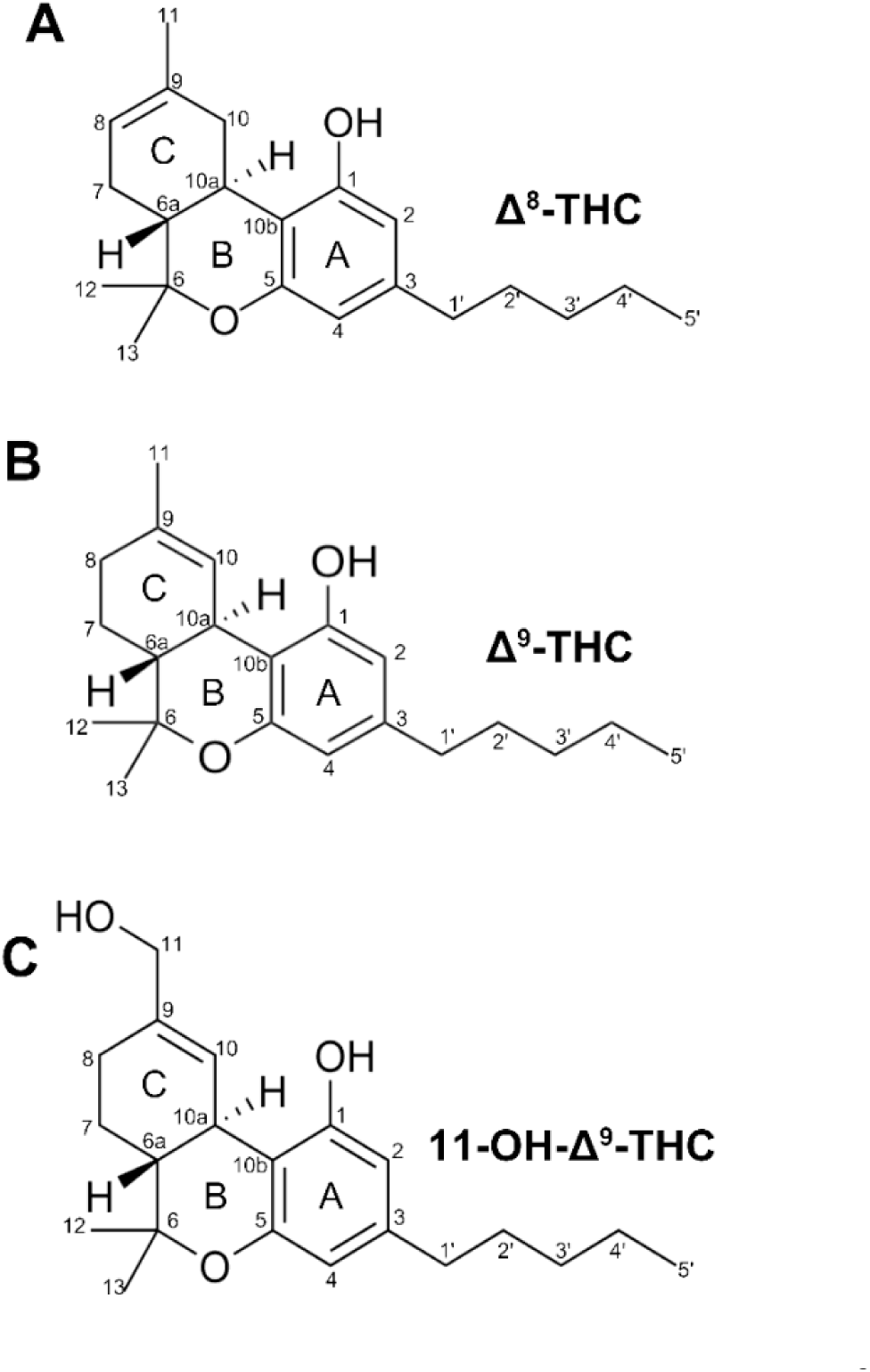
Chemical structures of (A) Δ^8^-tetrahydrocannabinol (Δ^8^-THC), (B) Δ^9^-tetrahydrocannabinol (Δ^9^-THC), and (C) 11-hydroxy-Δ⁹-tetrahydrocannabinol (11-OH-Δ^9^-THC), with IUPAC atom numbering shown. All three cannabinoids share the same tricyclic dibenzopyran scaffold (rings A, B, and C) and pentyl (aliphatic) side chain.

The exposure to Δ^8^-THC and Δ^9^-THC and the reported subjective pharmacodynamic effects were comparable when both compounds were taken via vaporization.^5^ In contrast, after oral dosing, THC exposure was higher but subjective pharmacodynamic effects were lower after Δ^8^-THC dosing than after Δ^9^-THC dosing.^6^ The differences in pharmacodynamic effects may be due to different formation kinetics of the psychoactive 11-OH-THC metabolite (**Figure 1**) from Δ^8^-THC and Δ^9^-THC and different pharmacological activity of 11-OH-Δ^8^-THC and 11-OH-Δ^9^-THC.

The psychoactive effects of cannabinoids are mainly mediated by cannabinoid receptor 1 (CB_1_) binding.^7^ Both Δ^8^-THC and Δ^9^-THC are partial agonists of CB_1_, and the potency of Δ^8^-THC appears overall weaker or similar to Δ^9^-THC across different *in vitro* assays of CB_1_ receptor binding.^8–10^ 11-OH-Δ⁹-THC shows ∼100-fold higher CB_1_ affinity than Δ^9^-THC at the human receptor but reduced efficacy in cyclic adenosine monophosphate (cAMP) inhibition.^11^ In rodent brain membranes, the affinity difference is modest.^12^ The CB_1_ binding affinity of 11-OH-Δ^8^-THC has not been directly compared to Δ^8^-THC, nor to 11-OH-Δ^9^-THC. The only currently available data compare 11-OH-Δ^8^-THC to Δ^9^-THC and show 11-OH-Δ^8^-THC inhibiting adenylate cyclase more potently than Δ^9^-THC in one study^10^ and less potently in another.^13^ As such, the extent to which 11-OH-Δ^8^-THC contributes to the effects of Δ^8^-THC remains unknown.

The pharmacological relevance of 11-OH-Δ^8^-THC is also impacted by exposure *in vivo*. Lower formation of 11-OH-THC has been proposed to explain the reduced *in vivo* potency of Δ^8^-THC.**^6^** After equal oral doses of Δ^8^-THC and Δ^9^-THC, 11-OH-Δ^8^-THC AUC was about 45% lower than 11-OH-Δ^9^-THC. Similar lower relative exposure of 11-OH-Δ^8^-THC was observed after vaporization.^5,6^ For Δ^9^-THC, the formation of 11-OH-THC is driven almost entirely by hepatic cytochrome P450 (CYP) 2C9, which is also the main clearance pathway of Δ^9^-THC in humans.^14^ Analysis of Δ^9^-THC depletion kinetics in human liver microsomes (HLM) resulted in an estimated CYP2C9 fraction metabolized (f_m_) of 0.91, with the residual f_m_ of 0.09 attributed to CYP2D6.^15^ Subsequently, systematic metabolite profiling in HLM found that Δ^9^-THC is oxidized to at least four main metabolites.^16^ The 11-OH-Δ^9^-THC formation was nearly entirely mediated by CYP2C9 but instead of CYP2D6, CYP3A was found to form the cyclohexenyl-ring hydroxylation products, and CYP2C19 contributed to the cyclohexenyl-ring hydroxylation and 11-hydroxylation.^16^ Further, minor metabolites were identified resulting from the hydroxylation of the pentyl side chain of Δ^9^-THC. Whether similar diversity in hydroxylation pathways exists for Δ^8^-THC is unknown.

Compared to Δ^9^-THC, the metabolism of Δ^8^-THC is poorly characterized. Recombinant CYP2C9 has been shown to form 11-OH-Δ^8^-THC from Δ^8^-THC at a lower catalytic rate than 11-OH-Δ^9^-THC from Δ^9^-THC.^17^ However, the enzymes contributing to hepatic Δ^8^-THC clearance have not been comprehensively defined, and the fractional contribution of 11-hydroxylation and CYP2C9 mediated metabolism to systemic Δ^8^-THC clearance remains unquantified. Based on the prior data that Δ^9^-THC is hydroxylated to at least four different mono-oxidation products in HLM by a variety of CYPs, we hypothesized that Δ^8^-THC, due to its structural similarity to Δ^9^-THC, also undergoes diversity of hydroxylation reactions by different CYPs in the human liver. Moving the double bond from the 9,10- to the 8,9-position alters the electronic environment of the C-ring carbons, while leaving the aliphatic side chain and the aromatic A-ring unchanged. We hypothesized that the allylic side chain and C11 hydroxylations of Δ^8^-THC are catalyzed similarly to Δ^9^-THC, while the cyclohexenyl ring hydroxylations would differ due to the position of the double bond. To test this hypothesis, the oxidation pathways of Δ^8^-THC were characterized in HLMs and in CYP Supersomes and the main enzymes contributing to the formation of Δ^8^-THC metabolites in humans were identified. We further aimed to define the role of CYP genetic polymorphisms in Δ^8^-THC metabolism and *in vivo* pharmacokinetics of Δ^8^-THC.

## 2. Materials and Methods

### 2.1. Chemicals, Reagents, Recombinant Enzymes and Human Liver Fractions

Cannabinoid reference standards Δ^8^-tetrahydrocannabinol (Δ^8^-THC; 1 mg/mL), Δ^9^-tetrahydrocannabinol (Δ^9^-THC; 1 mg/mL), Δ^9^-THC-d₃ (0.1 mg/mL), (±)-11-hydroxy-Δ^8^-THC (11-OH-Δ^8^-THC; 1 mg/mL), (±)-11-hydroxy-Δ^9^-THC (11-OH-Δ^9^-THC; 1 mg/mL), and (±)-11-OH-Δ^9^-THC-d₃ (0.1 mg/mL) were purchased as methanol solutions from Millipore-Sigma (St. Louis, MO). 8β,9β-epoxy-hexahydrocannabinol (Δ^8^-THC epoxide; 100 µg/mL) was purchased from Cayman Chemical Company (Ann Arbor, MI). β-nicotinamide adenine dinucleotide phosphate (NADPH) tetrasodium salt, dipotassium phosphate, monopotassium phosphate, potassium phosphate, dimethyl sulfoxide (DMSO), dextromethorphan, dextrorphan, dextrorphan-d₃, phenacetin, quinidine, tienilic acid, (+)-N-3-benzylnirvanol, troleandomycin, furafylline, and racemic fluoxetine were purchased from Millipore-Sigma (St. Louis, MO). Magnetic silica beads were obtained from Thermo Fisher Scientific (Waltham, MA). Acetonitrile, methanol, water, and formic acid used for liquid chromatography-tandem mass spectrometry (LC-MS/MS) were Optima grade and purchased from Thermo Fisher Scientific (Waltham, MA).

Recombinant human CYP1A1, CYP1A2, CYP1B1, CYP2B6, CYP2C8, CYP2C9, CYP2C19, CYP2D6, CYP2E1, CYP2J2, CYP3A4, CYP3A5, and CYP3A7 Supersomes co-expressed with cytochrome P450 reductase and cytochrome b_5_, were obtained from Corning Life Sciences (Woburn, MA). Human CYP2C9 variant Supersomes co-expressed with cytochrome P450 reductase (CYP2C9*1, cat. no. 456218, lot 27937; CYP2C9*2, cat. no. 456209, lot 2272872; CYP2C9*3, cat. no. 456242, lot 20439) were a gift from Discovery Life Sciences. Purified human P450 reductase was purchased from BD Gentest (Woburn, MA). Cytochrome C from equine heart (C7752, lot SLBQ2826V) was purchased from Millipore Sigma (St. Louis, MO, USA). Human liver microsomes (HLM) pooled from 50 individual donors were purchased from Gibco (Cat. No. HMMCPL; Thermo Fisher Scientific, Waltham, MA). Human liver microsomes from twenty-two previously described donors^16^ were obtained from the University of Washington (UW) Human Liver Bank. Donors ranged in age from 7 to 64 years and comprised of 7 females and 15 males (20 Caucasian, 1 Asian, 1 Black or African American).

### 2.2. General Incubation Protocol

Unless otherwise stated, all incubations were performed in Eppendorf low protein binding tubes in 100 mM potassium phosphate buffer pH 7.4 (KPi) at 37 °C in a shaking water bath. All incubations were conducted under conditions of time linearity established prior to kinetic experiments. Metabolite formation from Δ^8^-THC was evaluated in pooled human liver microsomes (HLM) and in a panel of recombinant human CYP enzymes. For incubations, HLM were diluted with KPi buffer to a final protein concentration of 0.1 mg/mL and recombinant CYP enzymes to 2 pmol/mL. Δ^8^-THC was added as methanol solution, and the final organic solvent content did not exceed 1% (v/v). Reaction mixtures with protein and Δ^8^-THC were pre-incubated for 5 minutes at 37 °C prior to initiating the reactions by the addition of 20 µL NADPH (final concentration 1 mM, total reaction volume, 200 µL). Recombinant CYPs were incubated for 30 minutes to allow detection of minor CYP activity and HLMs were incubated for 15 minutes. Reactions were terminated by the addition of 200 µL ice-cold acetonitrile containing 1% formic acid and 11-OH-Δ^9^-THC-d₃ (10 nM) as the internal standard. Samples were centrifuged at 18,000 × g for 30 minutes at 4 °C, and 150 µL of the resulting supernatant was transferred to glass inserts in autosampler vials for LC–MS/MS analysis. The dependence of metabolite formation on NADPH, HLM protein and substrate was confirmed by control incubations lacking either NADPH, protein or substrate. All incubations were performed in duplicate and repeated on at least three separate days unless otherwise noted.

The Supersomes containing polymorphic CYP2C9 were available co-expressed with P450 reductase, but reductase expression can differ between lots and genotypes. To address this potential confounding factor when comparing the genotypes, the reductase activity in the CYP2C9 *1, *2, and *3 Supersomes was quantified using a spectroscopic cytochrome c assay (**Supplemental Methods 1.1**). Purified human reductase was then added to all incubations with CYP2C9 *1, *2, and *3 Supersomes to achieve fixed reductase activity-to-CYP ratio (5 activity units of reductase per 1 nM CYP2C9) The incubations were then conducted as described above.

### 2.3. General Liquid Chromatography-Tandem Mass Spectrometry (LC-MS/MS) Method

Unless otherwise stated, all samples containing Δ^8^-THC and its metabolites were analyzed by LC-MS/MS using a Sciex API5500 QTRAP mass spectrometer (Framingham, MA) coupled to an Agilent 1290 Infinity Ultra-High-Performance Liquid Chromatography (UHPLC) system (Santa Clara, CA). Chromatographic separation was achieved using a Kinetex Evo C18 column (2.1 × 100 mm, 2.6 µm; Phenomenex) with the column oven set at 50 °C. Mobile phase consisted of water (A) and acetonitrile (B), both containing 0.1% formic acid. Gradient elution at a flow rate of 0.6 mL/min was used. The gradient was as follows: initial conditions 90% A and 10% B held for 1 minute, followed by a linear increase to 55% B at 5.0 minutes, 75% B at 8.0 minutes, and 90% B at 10.0 minutes, held at 90% B for 0.5 minutes before returning to initial conditions for 2 minutes prior to the next injection.

Analytes were detected in positive ion electrospray ionization (ESI) mode. Unless otherwise stated, instrument source parameters were as follows: ion spray voltage (IS) 5500 V, source temperature (TEM) 550 °C, curtain gas (CUR) 35 psi, ion source gas 1 (GS1) 60 psi, ion source gas 2 (GS2) 50 psi, and collision-activated dissociation (CAD) 2. Additional parameters included declustering potential (DP) 35 V, entrance potential (EP) 4 V, collision energy (CE) 33 V, collision cell exit potential (CXP) 10 V, and dwell time 100 ms per transition.

### 2.4. Identification of Δ^8^-THC Metabolites by LC-MS/MS

To identify Δ^8^-THC metabolites formed by CYPs, pooled human liver microsomes (HLM) and recombinant human CYP enzymes were incubated with Δ^8^-THC (5 µM final concentration) as described in **Section 2.2**. The incubations were initially analyzed by LC-MS/MS monitoring the MRM transitions *m/z* 331 → 201 and *m/z* 331 → 193 that were previously described^16^ for detection of Δ^9^-THC metabolites.

To obtain structural information on Δ^8^-THC metabolites, spectra were collected via enhanced product ion (EPI) scans in positive ion mode in samples from HLM and CYP Supersome incubations. MS/MS spectra of the [M + H]^+^ ions at *m/z* 331 were collected. Product ion spectra were acquired over a mass range of *m/z* 50–340. The EPI scan rate was set to 10,000 Da/s, dynamic fill time was enabled for the linear ion trap, Q1 was operated at unit resolution, and the Q3 entry barrier was set to 8 V. All LC and source parameters were identical to those used for MRM analyses described in section 2.3, except for collision energy (CE), which was varied stepwise from 10 to 50 V in increments of 10 V. The spectra shown for fragmentation analysis were collected with CE of 30 V. Following detection of metabolites formed and collection of metabolite spectra, dominant product ions were identified for each metabolite and used to select MRM transitions for quantification. The *m/z* 331 → 201 transition was used for 11-OH-Δ^8^-THC, M3, and M4 quantification, while the *m/z* 331 → 191 transition was used for M1 and M2 quantification.

### 2.5. Determination of Δ^8^-THC unbound fraction in microsomes and in recombinant CYP enzyme incubations

The unbound fraction of Δ^8^-THC in human liver microsomes (f_u,mic_) and Supersomes (f_u,CYP_) was determined in KPi using magnetic silica beads (MGSB), as previously described.^16^ In brief, for HLM 200 µL MGSB (21 x 10^9^ beads) and for recombinant CYPs 50 µL MGSB (5.25 x 10^9^ beads) were washed three times with 1 mL KPi buffer prior to use. Pooled HLMs (1.0 mg/mL in KPi), CYP1A2, CYP2C9, CYP2C19, CYP2D6, and CYP3A4 Supersomes (2 pmol/mL each), and polymorphic CYP2C9 Supersomes (1 pmol/mL each) were mixed with Δ^8^-THC at enzyme-specific nominal concentrations (1 and 2.5 µM for HLM, 2.5 µM for CYP1A2, 0.1 µM for CYP2C9 and its polymorphic variants, 0.3 µM for CYP2C19, 2 µM for CYP2D6, and 1 µM for CYP3A4), and pre-incubated for 5 min at 37°C in a shaking water bath to allow equilibration of Δ^8^-THC with protein. Following pre-incubation, 100 µL aliquots were removed as “Total” samples (no beads). The remaining mixture (450 µL) was transferred to tubes containing pre-washed magnetic silica beads and incubated for 30 min at 37 °C in a shaking water bath to maintain uniform bead suspension. After incubation, the beads were magnetized, and a 100 µL aliquot of the supernatant was collected to measure the free, unbound concentration. 200 µL of acetonitrile with 1% formic acid (v/v) and Δ^9^-THC-d_3_ internal standard were added to all samples, samples were centrifuged at 18,000 × g for 30 min at 4°C, and supernatants were transferred to MS vials for LC-MS/MS analysis. Δ^8^-THC was quantified by LC-MS/MS in positive ion electrospray ionization mode using multiple reaction monitoring (MRM). The MRM transitions used for Δ^8^-THC and Δ^8^-THC-d_3_ were *m/z* 315 > 193 m/z and *m/z* 318 > 196, respectively. Δ^8^-THC was quantified against two calibration curves (low curve, 0.8–300 nM; high curve, 47–5000 nM) prepared in KPi. Calibrators and incubation samples were processed identically, including a 10-fold dilution prior to analysis. Analytes were quantified using peak area ratios to the internal standard. The f_u,mic_ and f_u,CYP_ values were calculated as the ratio of the unbound to the total analyte concentration. All experiments were performed in duplicate and repeated on three different days. Values are reported as mean ± standard deviation.

### 2.6. Kinetics of Δ^8^-THC metabolite formation in pooled human liver microsomes and with recombinant CYPs

The kinetics of Δ^8^-THC metabolite formation were characterized in incubations performed within the time-linear range for metabolite formation with pooled HLMs and with recombinant CYP1A2, CYP2C9, CYP2C19, CYP2D6, and CYP3A4 Supersomes. The incubations were conducted as described in **Section 2.2**. Nominal substrate concentrations ranged from 0.05–15 µM for HLM and 0.05–5 µM for recombinant CYP incubations. 11-OH-Δ^8^-THC formation velocities were determined based on an authentic reference standard. Relative formation velocity was estimated for metabolites M1–M4 based on MRM peak area ratios between metabolite peak area and the internal standard peak area. All incubations were conducted in duplicate, and the kinetic experiments were repeated on three different days to ensure reproducibility.

To characterize 11-OH-Δ^8^-THC formation kinetics by polymorphic CYP2C9, Δ^8^-THC (10–640 nM nominal) was incubated with 1 nM recombinant CYP2C9 (*1, *2, or *3) reconstituted with cytochrome P450 reductase (**Supplemental Methods 1.1**). Reductase incorporation into the membrane was achieved as previously described^18^ by pre-incubating the purified reductase with the CYP2C9 Supersomes for 15 min at room temperature prior to adding substrate and KPi buffer to the final incubation volume. Reaction mixtures were pre-incubated for 5 min at 37 °C before initiating reactions with 1 mM NADPH, or buffer for no-NADPH negative controls. Reactions were allowed to proceed for 5 min at 37 °C before quenching with acetonitrile containing 1% formic acid and 11-OH-Δ⁹-THC-d_3_ internal standard. Quenched samples were centrifuged at 18,000 × g for 30 min prior to transferring the supernatant to vials for LC-MS/MS analysis.

Unbound Δ^8^-THC concentrations were derived prior to model fitting. The unbound fraction (f_u_) was measured for each protein preparation as described in **Section 2.5**, and nominal and total concentrations were then corrected for nonspecific binding to plastics and for microsomal protein binding using percent recovery values determined at multiple substrate concentrations relevant to the incubations. Unbound concentrations corresponding to each nominal concentration were calculated as [S]_u_ = f_u_ × [S]_nominal_. As shown before for Δ^9^-THC in Supersomes^16^ the f_u,mic_ for Δ^8^-THC in HLM did not differ between the two nominal concentrations tested, and the mean value was applied to all HLM incubations. Enzyme-specific f_u,CYP_ values were applied to the corresponding recombinant CYP incubations. All unbound concentrations were derived in this manner before any model was fit.

Metabolite formation velocities were plotted as a function of unbound Δ^8^-THC concentration, and kinetic parameters were estimated by nonlinear regression in GraphPad Prism (GraphPad Software, Boston, MA). This analysis was applied identically to HLM, recombinant CYP, and polymorphic CYP2C9 datasets, differing only in the normalization of velocity. For 11-OH-Δ^8^-THC formation in HLM, velocities were normalized to microsomal protein and the maximal velocity is reported as V_max_ (pmol min⁻¹ mg⁻¹ protein), whereas for recombinant CYP and polymorphic CYP2C9 incubations, velocities were normalized to CYP content and the maximal velocity is reported as the turnover number k_cat_ (pmol min⁻¹ nmol⁻¹ CYP). For metabolites M1–M4, maximum turnover is reported as peak area ratio (PAR) min⁻¹ mg⁻¹ protein or PAR min⁻¹ nmol⁻¹ CYP.

The Michaelis–Menten (**Equation 1**) and substrate inhibition (**Equation 2**) models were fit to data from both incubation systems. In Equations 1 and 2, the maximal rate term is V_max_ for pooled HLM incubations and the turnover number k_cat_ for recombinant CYP and polymorphic CYP2C9 incubations, in which velocities were normalized to CYP content. V_max_ is written below in the equations for clarity. Sigmoidal kinetics were observed only in pooled HLM incubations, and the Hill model (**Equation 3**) was therefore fit to HLM data only.

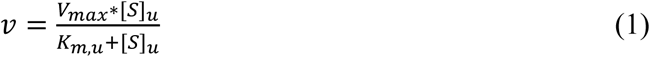

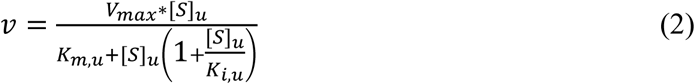

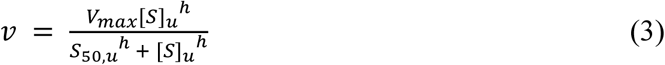

In equations 1-3, v is the reaction velocity, [S]_u_ is the unbound substrate concentration, V_max_ is the maximum reaction velocity, K_m,u_ is the unbound Michaelis constant, K_i,u_ is the unbound substrate inhibition constant, S_50,u_ is the unbound substrate concentration producing half-maximal velocity, and h is the Hill coefficient describing the degree of cooperativity.

Model selection was based on the corrected Akaike Information Criterion (AICc) computed in GraphPad Prism (GraphPad Software, Boston, MA), and the model yielding the lowest AICc was selected; the Hill model was included in this comparison for HLM datasets only. Because unbound concentrations were used as the independent variable throughout, the K_m,u_ and K_i,u_ estimated from the appropriate models are reported.

### 2.7. Prediction of the relative contribution of individual CYP enzymes to Δ^8^-THC metabolite formation

The contribution of individual hepatic CYP enzymes (f_m,CYP_) to specific Δ^8^-THC metabolite formation, was predicted from recombinant CYP activity data using literature-reported intersystem extrapolation factors (ISEFs) (**Supplemental Table S1**) and population-average hepatic CYP abundance values (**Supplemental Table S2**). The analyte to internal standard peak area ratio (PAR) was used as a surrogate for the formation rate of the metabolites M1-M4. The apparent intrinsic clearance (CL_app_) attributable to the specific CYP was estimated from **Equation 4**

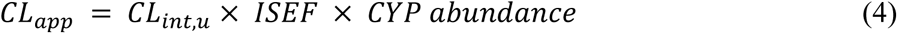

where CL_int,u_ is the unbound intrinsic clearance, ISEF is the intersystem extrapolation factor that scales intrinsic activity of the CYP between recombinant enzymes and the corresponding native enzyme in HLM, and CYP abundance is the hepatic protein abundance of each CYP (pmol min⁻¹ mg protein⁻¹). CL_app_ is expressed in µL min⁻¹ mg⁻¹ HLM protein. The fractional contribution of each CYP to the formation of a given metabolite was calculated from **Equation 5**

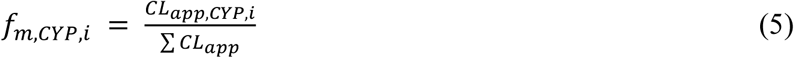

where f_m,CYP,i_ is the fractional contribution of CYP isoform i to the formation of a given metabolite, CL_app,CYP,i_ is the apparent intrinsic clearance attributable to that isoform (**Equation 4**), and ΣCL_app_ is the summed apparent intrinsic clearance across all CYP isoforms evaluated for the given metabolite.

### 2.8. Inhibition of Δ^8^-THC metabolism in pooled human liver microsomes

To estimate the relative contribution of individual CYP enzymes to specific Δ^8^-THC metabolite formation, Δ^8^-THC was incubated with pooled HLMs in the presence and absence of selective chemical inhibitors of CYP1A2 (furafylline), CYP2C9 (tienilic acid), CYP2C19 ((+)-N-3-benzylnirvanol), CYP2D6 (quinidine), and CYP3A (troleandomycin) and the multi-CYP inhibitor fluoxetine (CYP2D6, CYP2C19)^19^. With irreversible inhibitors (furafylline, 10 µM, tienilic acid, 50 µM, troleandomycin, 50 µM, and fluoxetine, 50 µM), HLMs (1 mg/mL in KPi) were incubated with the inhibitor (or with the corresponding vehicle; DMSO for furafylline, tienilic acid, and troleandomycin or methanol for fluoxetine) in the presence of 1 mM NADPH at 37°C for 30 min (tienilic acid^20^ and fluoxetine) or 10 min (furafylline^21^ and troleandomycin^21^). Following the inactivation step, the HLM mixture was diluted 20-fold in KPi buffer containing 1 mM NADPH to achieve a final protein concentration of 0.05 mg/mL, and reactions were initiated by the addition of Δ^8^-THC (2 µM or 0.3 µM final concentration). After incubation for 5 min at 37°C, reactions were quenched by adding acetonitrile with internal standard, and the samples were analyzed as described in **Section 2.3 and 2.4** to measure Δ^8^-THC metabolite formation.

For reversible inhibitors, inhibitor ((+)-N-3-benzylnirvanol 5 µM and quinidine 4 µM) or vehicle (methanol), HLMs (0.05 mg/mL in KPi), and Δ^8^-THC (2 µM or 0.3 µM final concentration) were mixed and preincubated at 37 °C for 5 min prior to initiating the reactions with addition of NADPH (1 mM). After 5 min ((+)-N-3-benzylnirvanol inhibition) or 15 min (quinidine inhibition),^21,22^ incubation, the reactions were quenched by adding acetonitrile with internal standard and processed as described in **Section 2.3 and 2.4**. All inhibition experiments were performed in duplicate and repeated on three independent experimental days. The f_m_ of each CYP isoform to Δ^8^-THC metabolite formation was estimated by comparing metabolite formation rates in inhibitor-treated versus vehicle control incubations.

### 2.9. Characterization of CYP1A2, CYP2C9, CYP2C19, CYP2D6 and CYP3A activity in individual donor human liver microsomes

The activities of CYP2C9, CYP2C19, and CYP3A, in the panel of 22 HLM donors used in this study were previously measured^16^ using tolbutamide (TBU; 10 µM), (S)-mephenytoin (MPH; 6 µM) and midazolam (MDZ; 1 µM) as probe substrates, respectively. CYP1A2 and CYP2D6 activities in the same HLM panel were measured in this study using phenacetin (5 µM) and dextromethorphan (1 µM) as selective probe substrates, respectively. Substrate concentrations were selected to be lower than 0.2 times reported K_m_ values.

The HLM incubations to measure acetaminophen formation from phenacetin and dextrorphan formation from dextromethorphan were conducted in a total reaction volume of 100 µL in 96-well plates as described in **Section 2.2**. After a 30 min incubation with phenacetin (5 µM) and 5 min with dextromethorphan (2 µM), reactions were quenched with 100 µL ice-cold acetonitrile containing 1% formic acid and 10 nM of the appropriate internal standard (acetaminophen-d_4_ or dextrorphan-d_3_), centrifuged (3,200 × g, 60 min), and analyzed by LC-MS/MS as described in **Supplemental Method 1.2**. Calibration curves ranged from 1.95 to 1,000 nM for acetaminophen and from 1.56 to 400 nM for dextrorphan. All incubations were performed within the time-linear range for metabolite formation and in technical duplicates repeated on three different days.

### 2.10. Correlation of Δ^8^-THC metabolism with CYP-specific activities

Δ^8^-THC incubations with the panel of 22 individual HLM donors were conducted as described in **Section 2.2**, using an incubation time of 5 min. This duration was selected because it fell within the linear range of metabolite formation while providing sufficient signal to detect the minor metabolites M1–M4. Simple linear regression was then used to test for correlation between specific CYP activities (CYP1A2, CYP2C9, CYP2C19, CYP2D6, and CYP3A) and the formation of Δ^8^-THC metabolites (11-OH-Δ^8^-THC, M1, M2, M3, and M4) across the panel. Due to the potential contributions by multiple enzymes to each metabolite formation, multiple linear regression models were subsequently constructed to test predictions by all relevant CYP activities as previously described.^16^ In the multiple linear regression models, the specific CYP activity measured using the probe substrates that had the lowest p-value in the simple linear regression model was selected as the base model for each metabolite. Relevant individual CYP activity was then added to the base model for each metabolite. To determine whether inclusion of additional CYP activities significantly improved model fit, the nested models were compared to the base model using F-tests with p-values < 0.01 considered statistically significant. All statistical analyses were performed using Python (version 3.9.6).

### 2.11. Impact of CYP genotypes on Δ^8^-THC metabolism in the human liver donors

The livers in the UW human liver bank have been previously genotyped for CYP2C9, CYP2C19, and CYP2D6. CYP2D6 diplotypes were assigned from PGRNseq next-generation sequencing data coupled with Stargazer structural-variant analysis.^23^ CYP2C9 diplotypes were determined from the same PGRNseq resequencing dataset.^24^ CYP2C19 diplotypes were from genotyping of the CYP2C19*2 (rs4244285) and *17 (rs12248560) variants together with CYP2C19 exon sequencing.^25^ Based on the genotypes, each of the 22 donors was assigned an activity score for CYP2C9, CYP2C19, and CYP2D6. For CYP2C9, allele functional values and the diplotype activity score (sum of the two allele values) followed the CPIC CYP2C9 activity-score system (*1 = 1.0, *2 = 0.5, *3 = 0).^26^ For CYP2C19, alleles were assigned numeric values consistent with their PharmVar/CPIC functional classifications (*17 = 1.5 (increased function), *2 = 0 (no function), *1 = 1.0 (normal)),^27^ and the diplotype activity score was calculated as the sum of the two allele values. For CYP2D6, each allele was assigned a functional value according to its CPIC function category (normal function = 1.0, decreased function = 0.5, no function = 0), and the diplotype activity score was calculated as the sum of the two allele values following the CPIC/DPWG consensus genotype-to-phenotype translation.^28–30^ Allele-to-function assignments and the resulting activity scores are provided in **Supplemental Table S3**.

The relationship between genotype-predicted CYP activity score and Δ^8^-THC metabolite formation was evaluated by Spearman rank correlation across the 22 donors. For each CYP, the donor activity score was correlated with the formation of 11-OH-Δ^8^-THC (pmol min⁻¹ mg⁻¹ protein) and of metabolites M1–M4 (peak area ratio per min per mg protein). Spearman correlation coefficients (ρ) and associated p-values were determined for each enzyme–metabolite pair using the cor.test function (stats package) in R (version 4.5.1; R Core Team, 2025), with statistical significance defined as p < 0.05. Data processing was performed using readxl, dplyr, and tidyr, and figures were generated using ggplot2, ggrepel, and patchwork.

### 2.12. Prediction of the effect of CYP2C9 genotype on 11-OH-Δ^8^-THC formation clearance

The kinetics of 11-OH-Δ^8^-THC formation by recombinant CYP2C9*1, *2, and *3 were characterized as described in **Section 2.6**. To predict the effect of CYP2C9 genotype on the relative contribution of CYP2C9 and CYP2C19 to 11-OH-Δ^8^-THC formation, the CYP2C9 CL_int,u_ term in **Equation 4** was replaced with the corresponding genotype-specific CL_int,u_ value, and f_m,CYP_ was recalculated using **Equation 5** for each of the six CYP2C9 diplotypes. Based on the data obtained with recombinant CYP2C9*1, *2, and *3 activity, the allele-specific activity was expressed relative to *1 as the ratio of CL_int,u_ to that of the *1 allele, giving 100% for *1/*1, 84% for *2/*2, and 6.0% for *3/*3. For heterozygous diplotypes, activity was assigned as the arithmetic mean of the two corresponding homozygous values, giving 92% for *1/*2, 53% for *1/*3, and 45% for *2/*3.

To predict the impact of substrate concentration and CYP2C9 genotypes on Δ^8^-THC CL_int,u_, CL_int,u_ of 11-OH-Δ^8^-THC in the human liver was predicted according to the Michaelis–Menten equation and substrate inhibition model (**Equations 1 and 2**) according to **Equation 6**:

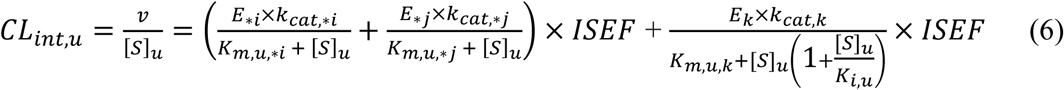

*E*_∗*i*_ and *E*_∗*j*_ represent the expression level of CYP2C9 proteoforms (*1, *2, or *3). In CYP2C9 heterozygous carriers each proteoform (*i,j*) was assumed to constitute 50% of the total CYP2C9 expression. The k_cat_ and K_m,u_ associated with each proteoform (*1, *2, or *3) acquired for CYP2C9 *1, *2, or *3 Supersomes were used for the prediction, expressed as k_cat,*i,_ k_cat,*j,_ K_m,u,*i,_ and K_m,u,*j_. *E_k_* represents the expression level of CYP2C19 protein. The k_cat_, K_m,u_, and K_i,u_ for CYP2C19 were used for the prediction, expressed as k_cat,k,_ K_m,u,k,_ and K_i,u_. To generate a simulation of 11-OH-Δ^8^-THC intrinsic formation clearance per mg microsomal protein as a function of the unbound Δ^8^-THC concentration, [*S*]*_u_* (0.1 *nM* − 100 *nM*) was swept. The Δ^8^-THC concentration was chosen to bracket circulating concentrations reported after oral and vaporized administration. The fraction of 11-OH-Δ^8^-THC metabolized by CYP2C9 or CYP2C19 was simulated using **Equation 5**, where the total clearance was calculated using **Equation 6**.

## 3. Results

### 3.1. Identification of Δ^8^-THC metabolites formed by cytochrome P450 enzymes

Several NADPH dependent oxidation products of Δ^8^-THC were detected by LC–MS/MS in incubations of Δ^8^-THC with HLM (**Figure 2**). The protonated molecular ion [M+H]⁺ of *m/z* 331 is consistent with the addition of a single oxygen atom to the parent cannabinoid (MW = 314). Extracted ion chromatograms of the diagnostic MRM transitions *m/z* 331→193 and *m/z* 331→201 showed five metabolite peaks that eluted between 5.9 and 6.4 min. These peaks were designated 11-OH-Δ^8^-THC and M1–M4 (**Figure 2**).

**Figure 2:**
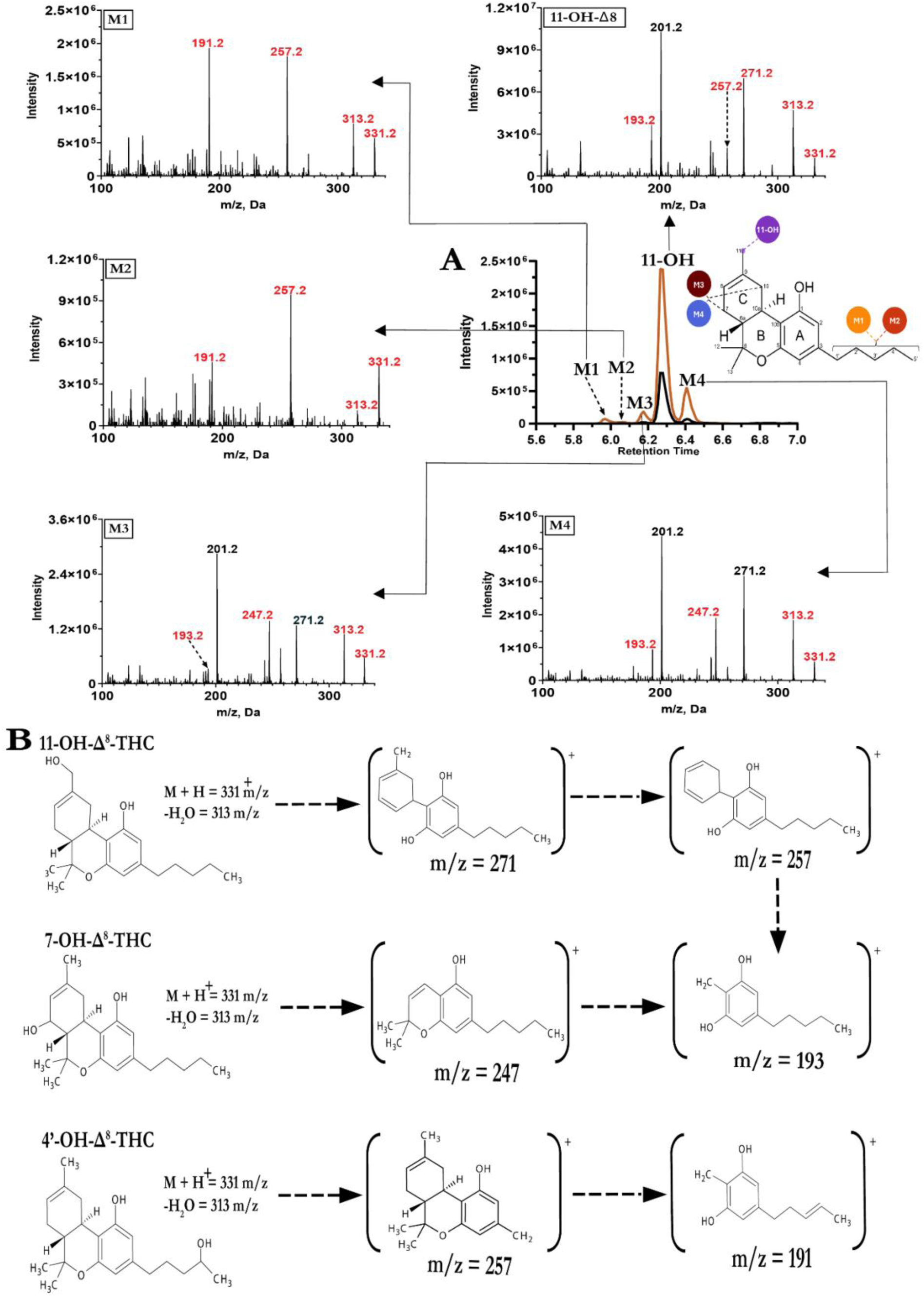
LC–MS/MS characterization of the monohydroxylated metabolites of Δ^8^-THC. (**A**) Extracted ion chromatograms (EICs) of the MS/MS transitions *m/z* 331→193 (black) and *m/z* 331→201 (gold) from incubations of Δ^8^-THC with human liver microsomes (HLM) are shown in the center, with the MS/MS spectra of the protonated molecular ion ([M+H]⁺, *m/z* 331.2) of each of the five metabolites arranged around the chromatogram and connected by arrows to the corresponding peak. M1 and M2 display the *m/z* 191.2 fragment diagnostic of aliphatic side-chain hydroxylation; M3 and M4 display the *m/z* 247.2 and 193.2 fragments characteristic of C-ring hydroxylation. 11-OH-Δ^8^-THC displays *m/z* 271.2, 257.2 and 193.2. Proposed hydroxylation sites are indicated on the Δ^8^-THC structure (inset): 11-OH (purple), the C-ring positions (7 or 10) assigned to M3 (burgundy) and M4 (blue), and the aliphatic side-chain positions assigned to M1 (orange) and M2 (red). (**B**) Proposed fragmentation pathways and fragment ion structures for 11-OH-Δ^8^-THC, 7-OH-Δ^8^-THC (a representative C-ring hydroxylation product), and 4′-OH-Δ^8^-THC (a representative aliphatic side-chain hydroxylation product). All three pathways proceed via loss of water from the protonated molecular ion (*m/z* 331 → 313).

MS/MS spectra of the protonated molecular ion [M+H]⁺ of *m/z* 331 were collected for all five metabolites (**Figure 2A**). A fragment ion at *m/z* 313 was observed in all spectra, corresponding to neutral loss of water (−18 Da). The identity of 11-OH-Δ^8^-THC was confirmed by co-chromatography and spectral matching with a commercially available reference standard (**Supplemental Figure S1**). The 11-OH-Δ^8^-THC metabolite MS/MS spectrum (**Figure 2A**) showed prominent ions at *m/z* 271, *m/z* 257, *m/z* 201, and *m/z* 193, similar to the reference compound (**Supplemental Figure S1**), consistent with the proposed fragmentation pathway (**Figure 2B**) and similar to previously reported^16^ 11-OH-Δ^9^-THC fragmentation.

The MS/MS spectra of M1 and M2 (**Figure 2A**) were both dominated by ions at *m/z* 257 and *m/z* 191, without notable *m/z* 201 or *m/z* 193 fragments. The presence of *m/z* 191 and absence of *m/z* 193 is consistent with the previously proposed^16^ fragmentation pattern for pentyl side-chain hydroxylated metabolites of Δ^9^-THC, and matches the proposed fragmentation pathway for 4′-OH-Δ^8^-THC (or other side chain hydroxylation products) shown in **Figure 2B**. M1 and M2 were therefore tentatively identified as pentyl side chain (1’-4’) hydroxylated metabolites of Δ^8^-THC (**Figure 2**).

The MS/MS spectra of M3 and M4 (**Figure 2A**) contained prominent ions at *m/z* 271, *m/z* 247 and *m/z* 201, with only a minor fragment at *m/z* 193, consistent with hydroxylation in the C-ring of Δ^8^-THC and matching the proposed fragmentation pathway for 7-OH-Δ^8^-THC shown in **Figure 2B**. M3 and M4 were therefore tentatively assigned as 7-or 10-OH-Δ^8^-THC (**Figure 2**). The formation of an 8,9-epoxide was also considered as a potential oxidation product of Δ^8^-THC. The synthetic 8,9-epoxide-Δ^8^-THC eluted later than any of the metabolites formed in HLM and had a different MS/MS spectrum (**Supplemental Figure S1**), with *m/z* 259 and *m/z* 231 fragments. These fragments were not observed in the spectra of M3 or M4, ruling out the possibility that M3 or M4 is the epoxide metabolite.

### 3.2. Kinetics of Δ^8^-THC metabolism in human liver microsomes

The kinetics of M1, M2, M3, M4, and 11-OH-Δ^8^-THC formation were characterized as a function of unbound Δ^8^-THC concentration in HLM. The f_u_ of Δ^8^-THC in HLM was 0.024 ± 0.003 and independent of Δ^8^-THC concentration. Due to the high nonspecific binding of Δ^8^-THC, all nominal concentrations were corrected for nonspecific adsorption to plastics and the unbound fraction 0.024 was applied to the measured concentration in solution. Because authentic reference standards were not available for M1–M4, absolute V_max_ could not be determined for them. Formation of these metabolites was instead measured as a relative response (PAR min⁻¹ mg⁻¹ protein). Linearity of the peak area response for these metabolites was confirmed via dilution of an incubation sample generated at a saturating concentration of Δ^8^-THC.

Formation of M1 and M2 followed Michaelis-Menten kinetics, with Δ^8^-THC K_m,u_ values of 120 ± 9 nM and 86 ± 17 nM, respectively (**Figure 3A, 3B**). Formation of M3 followed sigmoidal kinetics, yielding an S_50,u_ of 46 ± 3 nM and a Hill coefficient (h) of 1.75 ± 0.08 (**Figure 3C**). Formation of 11-OH-Δ^8^-THC followed substrate inhibition kinetics, with a K_m,u_ of 3.6 ± 0.2 nM and a V_max_ of 325.0 ± 6.9 pmol min⁻¹ mg⁻¹ protein (**Figure 3D**). Substrate inhibition was weak over the concentration range tested, and the K_i,u_ could only be estimated as >910 nM. Formation of M4 followed Michaelis-Menten kinetics but the 95% confidence interval of the fitted K_m,u_ value exceeded the concentration range of Δ^8^-THC tested in the incubations. Hence, only the lower bound of the K_m,u_ (121 nM) is reported (**Figure 3E**). The K_m,u_ for 11-OH-Δ^8^-THC formation was more than an order of magnitude lower than that for any other metabolite. Collectively, the differences in the K_m,u_ values of Δ^8^-THC metabolite formation suggest that multiple enzymes contribute to Δ^8^-THC metabolism in HLM.

**Figure 3:**
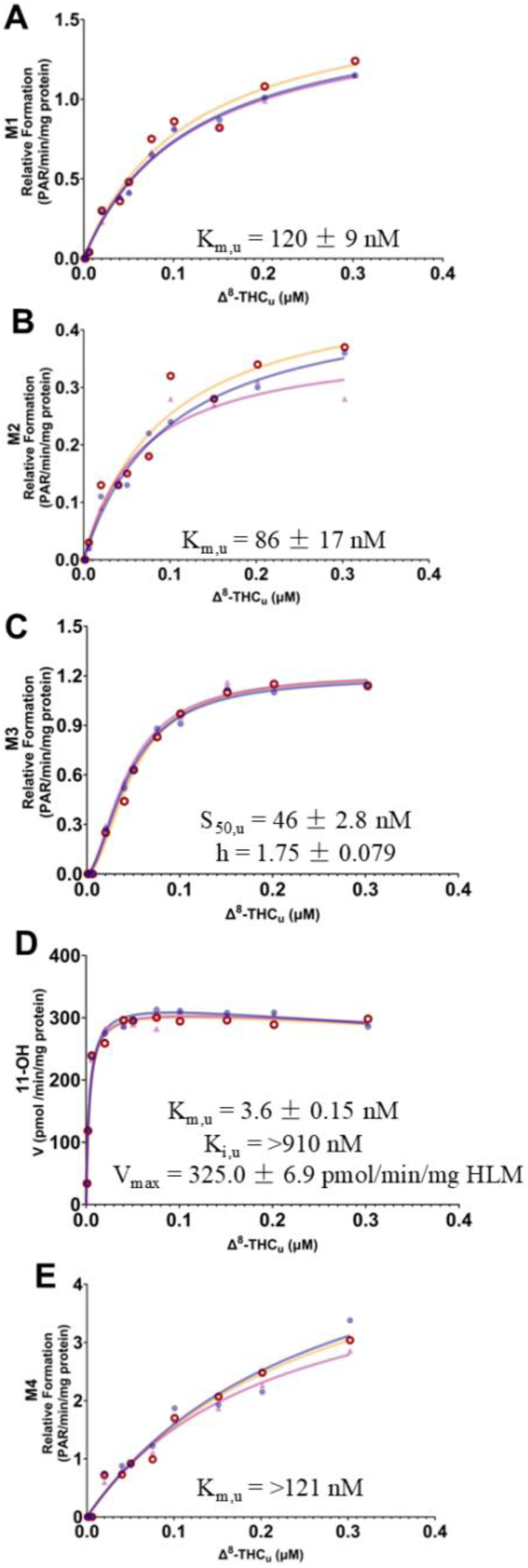
Kinetics of Δ^8^-tetrahydrocannabinol (Δ^8^-THC) metabolite formation in human liver microsomes (HLM). Formation of (**A**) M1, (**B**) M2, (**C**) M3, (**D**) 11-hydroxy-Δ^8^-THC (11-OH-Δ^8^-THC), and (**E**) M4 is plotted against unbound Δ^8^-THC concentration. M1–M4 are expressed as relative formation (peak area ratio (PAR) min⁻¹ mg⁻¹ protein) because authentic reference standards were unavailable where 11-OH-Δ^8^-THC is expressed as velocity (pmol min⁻¹ mg⁻¹ protein). Data are from *n* = 3 independent experiments, each shown as a separate symbol and color. Solid lines are best fits to the Michaelis-Menten equation (**A, B, E**), the Hill equation (**C**), or a substrate inhibition model (**D**). Kinetic parameters derived from these fits are shown as insets and are reported as mean ± standard deviation across replicate experiments. For M4 (**E**), formation did not approach saturation over the concentration range tested and the K_m,u_ shown is therefore a minimum estimate rather than a precisely determined value.

### 3.3 Identification of cytochrome P450 enzymes that metabolize Δ^8^-THC and predictions of relative contributions of the CYPs to metabolite formation

To identify the CYP enzymes responsible for the formation of Δ^8^-THC metabolites in HLMs, Δ^8^-THC was incubated with a panel of thirteen recombinant CYPs (**Figure 4**), and the metabolite formation was monitored using the most abundant MRM transition for each metabolite (**Supplemental Figure S2**). Of the enzymes tested, CYP2C19 had the highest relative activity towards 11-OH-Δ^8^-THC formation, followed by CYP2C9 (**Figure 4**). Modest 11-OH-Δ^8^-THC formation was also detected with CYP2B6 and CYP2D6 (**Figure 4**). M1 formation was most abundant in incubations with CYP2D6, with lower formation observed with CYP1A1, CYP1A2, CYP2J2, CYP3A4, and CYP3A5 (**Figure 4**). M2 formation was detected with CYP1A1, CYP1A2, CYP2C19, CYP2D6, CYP2J2, CYP3A4, and CYP3A5 (**Figure 4**). M3 and M4 were only formed by the CYP3A enzymes.

**Figure 4:**
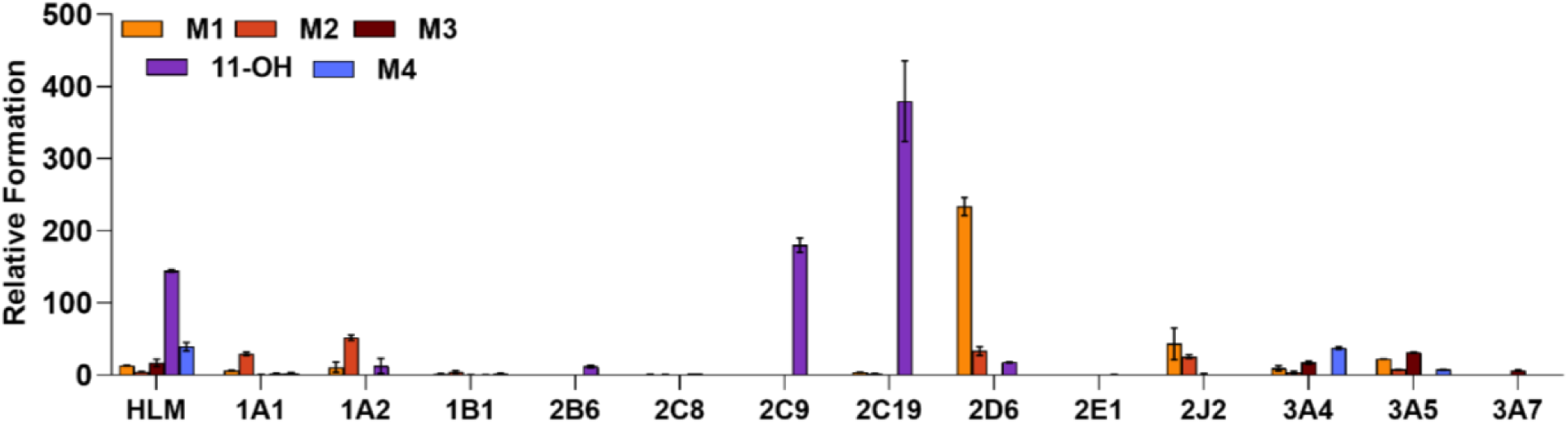
Relative formation of Δ^8^-THC metabolites in pooled human liver microsomes and in a panel of recombinant CYP enzymes. Relative formation of monohydroxylated metabolites M1–M4 and 11-OH-Δ^8^-THC in incubations of Δ^8^-THC (5 µM) with pooled HLM or a panel of recombinant CYP enzymes (10 pmol/mL) in the presence of NADPH. Each metabolite was quantified using the most abundant MRM transition: M1 and M2 by *m/z* 331→191 and M3, 11-OH-Δ^8^-THC, and M4 by 331→201. Metabolite formation is expressed relative to the internal standard 11-OH-Δ^9^-THC-d₃. The data are a representative experiment performed in triplicate.

Based on the identification of candidate CYPs contributing to Δ^8^-THC metabolism in HLM, the kinetics of Δ^8^-THC metabolite formation were characterized using recombinant CYP1A2, CYP2C9, CYP2C19, CYP2D6, and CYP3A4 (**Figure 5A-E**). The kinetic parameter estimates are summarized in **Table 1**. The formation of all metabolites by the tested CYPs, except 11-OH-Δ^8^-THC formation by CYP2C19, followed Michaelis-Menten kinetics (**Figure 5**, **Table 1**). 11-OH-Δ^8^-THC formation by CYP2C19 was best described by a substrate inhibition model. M2 formation by CYP2C19 was too low for reliable quantification. For 11-OH-Δ^8^-THC formation by CYP2D6 and M4 formation by CYP3A4, the 95% confidence interval of the fitted K_m,u_ value exceeded the concentration range of Δ^8^-THC in the incubations. Hence, K_m,u_ is reported as the lower bound of the 95% CI for K_m,u_ and k_cat_ is not reported for these reactions.

**Figure 5:**
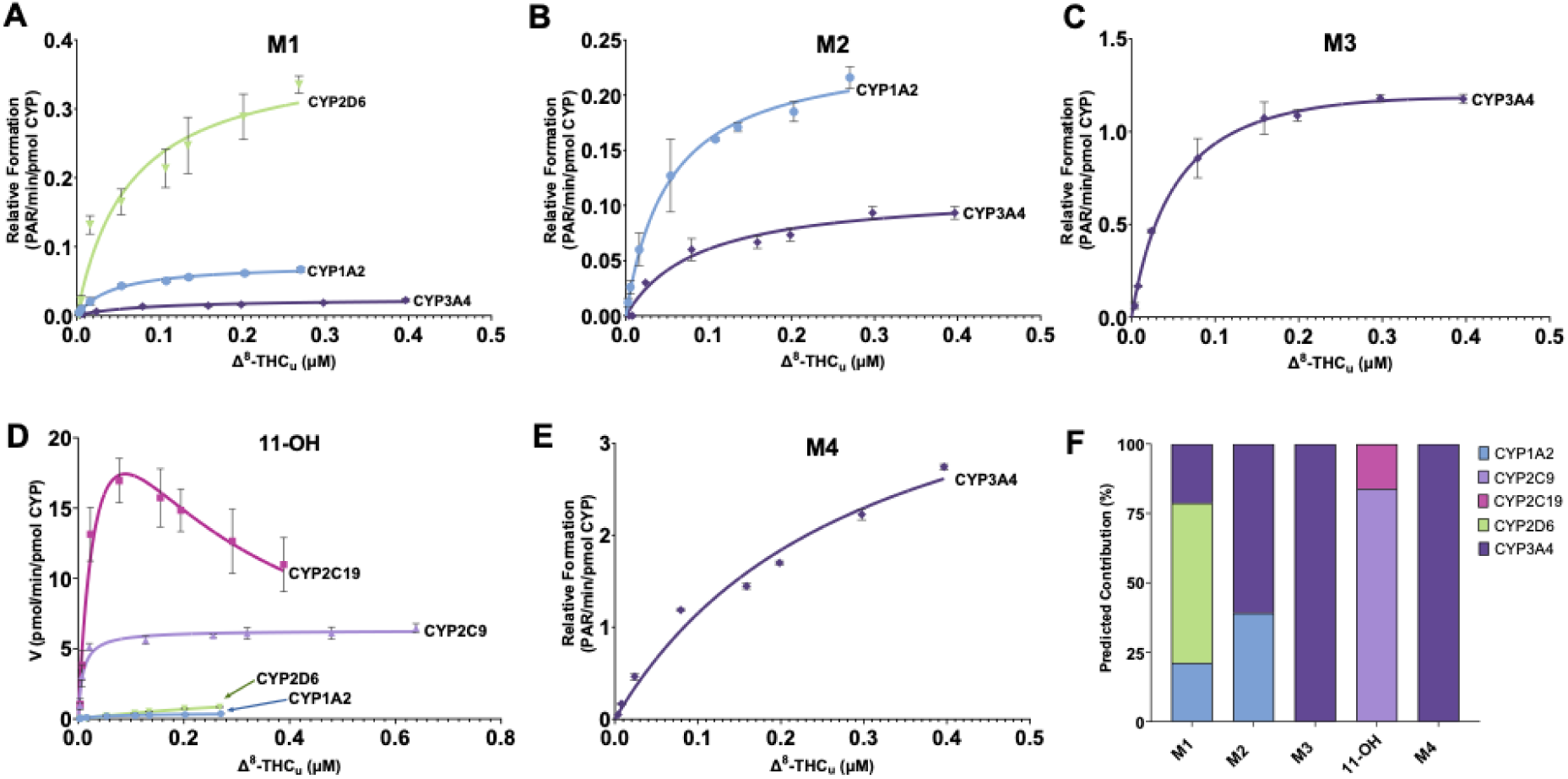
Kinetics of Δ^8^-THC metabolite formation by recombinant CYPs. (**A**) M1 formation by CYP2D6 (green), CYP1A2 (light blue), and CYP3A4 (purple). (**B**) M2 formation by CYP1A2 and CYP3A4. (**C**) M3 formation by CYP3A4. (**D**) 11-OH-Δ^8^-THC formation by CYP2C19 (purple), CYP2C9 (blue), CYP2D6 (green), and CYP1A2 (light blue). (**E**) M4 formation by CYP3A4. (**F**) Predicted fractional contribution (%) of each CYP to the formation of the individual metabolites M1–M4 and 11-OH-Δ^8^-THC based on the enzyme kinetics as described in **Section 2.7** (CYP1A2 – light blue, CYP2C9 – light purple, CYP2C19 – pink, CYP2D6 – green, CYP3A4 – dark purple). Data were fit to the Michaelis–Menten equation or the substrate inhibition equation, as appropriate for each enzyme– metabolite pair. For M1–M4, formation is expressed as relative formation (peak area ratio; PAR min⁻¹ pmol⁻¹ CYP), while 11-OH-Δ^8^-THC is expressed as pmol min⁻¹ pmol⁻¹ CYP.

**Table 1:** Kinetic parameters for the formation of Δ^8^-THC metabolites M1–M4 and 11-OH-Δ^8^-THC by recombinant human CYP enzymes. Parameters were derived by nonlinear regression of metabolite formation rates versus unbound Δ^8^-THC concentration using the Michaelis–Menten, or substrate inhibition models, as appropriate. K_m_,_u_ Michaelis constant (Michaelis–Menten); K_i,u_, unbound substrate inhibition constant; k_cat_, catalytic rate constant; CL_int_, intrinsic clearance, calculated as k_cat_/K_m,u_. Values are the parameter estimates ± SD from duplicate experiments conducted on 3 individual days. Absolute k_cat_ could be determined only for 11-OH–Δ^8^-THC, for which an authentic standard was available; for M1–M4, formation was measured as relative peak area ratio. N.D., not determined.

| CYP | Substrate | $K_{m,u}$ (nM) | $k_{cat}^a$ | $CL_{int}^a$ |
| --- | --- | --- | --- | --- |
| **CYP1A2** | $f_u = 0.059 \pm 0.0046$ | | | |
| | M1 | $44 \pm 6$ | $74 \pm 4.2$ | $1.7 \pm 0.4$ |
| | M2 | $52 \pm 35$ | $240 \pm 9.6$ | $4.9 \pm 1.8$ |
| | 11-OH-$\Delta^8$-THC | $61 \pm 23$ | $410 \pm 75$ | $7.4 \pm 3.1$ |
| **CYP2C9** | $f_u = 0.17 \pm 0.029$ | | | |
| | 11-OH-$\Delta^8$-THC | $8.8 \pm 2$ | $6300 \pm 0.19$ | $716 \pm 90$ |
| **CYP2C19** | $f_u = 0.096 \pm 0.019$ | | | |
| | 11-OH-$\Delta^8$-THC<sup>b</sup> | $40 \pm 5$ | $35310 \pm 1.36$ | $882 \pm 68$ |
| **CYP2D6** | $f_u = 0.080 \pm 0.0065$ | | | |
| | M1 | $64 \pm 19$ | $380 \pm 0.03$ | $6.0 \pm 1.7$ |
| | 11-OH-$\Delta^9$-THC | $>346^c$ | N.D. | N.D. |
| **CYP3A4** | $f_u = 0.10 \pm 0.0094$ | | | |
| | M1 | $84 \pm 11$ | $25 \pm 0.6$ | $0.30 \pm 0.05$ |
| | M2 | $87 \pm 20$ | $110 \pm 6.9$ | $1.3 \pm 0.4$ |
| | M3 | $47 \pm 2.1$ | $1350 \pm 35$ | $28.7 \pm 2.1$ |
| | M4 | $>135^c$ | N.D. | N.D. |

Values are mean ± SD. The unbound fraction ( $f_u$ ) is enzyme-specific and is given in each enzyme heading row, and applies to all data within that block.
<sup>a</sup> $k_{cat}$ for 11-OH- $\Delta^8$ -THC is expressed as $\text{pmol min}^{-1} \text{nmol}^{-1}$ CYP. $k_{cat}$ for M1–M4 is expressed as peak area ratio (PAR) $\text{min}^{-1} \text{nmol}^{-1}$ CYP as authentic standards were unavailable for absolute quantification.. $CL_{int}$ is expressed using units corresponding to $k_{cat}$ .
<sup>b</sup>Substrate inhibition was observed for CYP2C19 only; $K_{i,u} = 170 \pm 15$ nM. No substrate inhibition was detected for any other enzyme–substrate pair.
<sup>c</sup>Formation did not reach saturation over the concentration range tested, and $K_{m,u}$ could not be estimated with precision. Therefore, the upper bound of the 95% confidence interval exceeded the highest concentration tested. The value shown is the lower bound of the 95% confidence interval and is therefore reported as a minimum estimate. Because $K_{m,u}$ was not defined, $k_{cat}$ could not be determined for these reactions. The highest unbound concentrations tested were 268 nM for CYP2D6 and 397 nM for CYP3A4.

The f_m_ values for each Δ^8^-THC metabolite by specific enzymes in human liver lacking CYP3A5 expression were predicted from unbound intrinsic clearance (k_cat_/K_m,u_) scaled to hepatic CYP abundance after incorporating ISEF values (**Figure 5F**). Because this approach assumes intrinsic clearances and S_u_<< K_m,u_, the predictions apply to unbound Δ^8^-THC concentrations of <3 nM for 11-OH-Δ^8^-THC and <20 nM for other metabolites. M1 formation was predicted to be mediated mainly by CYP2D6 (f_m_ = 0.58), with smaller and comparable contributions from CYP1A2 (f_m_ = 0.21) and CYP3A4 (f_m_ = 0.21). M2 formation was predicted to be mediated largely by CYP3A4 (f_m_ = 0.60), with the remainder attributable to CYP1A2 (f_m_ = 0.40). Despite CYP1A2 having the higher unbound intrinsic clearance of the two enzymes, the lower hepatic abundance of CYP1A2 relative to CYP3A4 resulted in the higher predicted contribution by CYP3A4. CYP2C9 was predicted to be the primary contributor to 11-OH-Δ^8^-THC formation (f_m_ = 0.84), followed by CYP2C19 (f_m_ = 0.16), with negligible contributions from CYP1A2 and CYP2D6. M3 and M4 were predicted to be formed exclusively by CYP3A4 (f_m_ = 1.0 for each).

### 3.3. Characterization of the CYP enzymes contributing to Δ^8^-THC hydroxylation in human liver microsomes

To test whether the predicted relative contributions of CYP enzymes to formation of specific Δ^8^-THC metabolites translate to HLMs, chemical inhibition experiments were conducted in pooled HLMs using selective inhibitors of CYP1A2, CYP2C9, CYP2C19, CYP2D6, and CYP3A, together with the multi-CYP inhibitor fluoxetine (**Figure 6**). Two different total concentrations of Δ^8^-THC, 2 µM and 0.3 µM corresponding to unbound concentrations of 40 nM and 6 nM, were used in the inhibition experiments. The CYP2C9 irreversible inhibitor tienilic acid inhibited 11-OH-Δ^8^-THC formation by ∼85%, indicating that CYP2C9 is the main enzyme forming 11-OH-Δ^8^-THC (**Figure 6**). Inhibitors of CYP3A, CYP2D6, or CYP1A2 did not decrease 11-OH-Δ^8^-THC formation. Despite recombinant CYP2C19 having a relatively high CL_int_ for 11-OH-Δ^8^-THC formation, benzylnirvanol did not inhibit 11-OH-Δ^8^-THC formation in HLM at the higher substrate concentration but modest (<10%) inhibition was observed at the lower substrate concentration. Competition between substrate and inhibitor may in part account for the different % inhibition at the two substrate concentrations. The unbound Δ^8^-THC concentrations were equal to (40 nM) or below (6 nM) the K_m,u_ of Δ^8^-THC for CYP2C19 (40 ± 5 nM, Table 1) in these experiments. Fluoxetine, an irreversible inhibitor of CYP2C19, reduced 11-OH-Δ^8^-THC formation modestly (∼10%) at both substrate concentrations, suggesting that CYP2C19 has a minor contribution to 11-OH-Δ^8^-THC formation.

**Figure 6:**
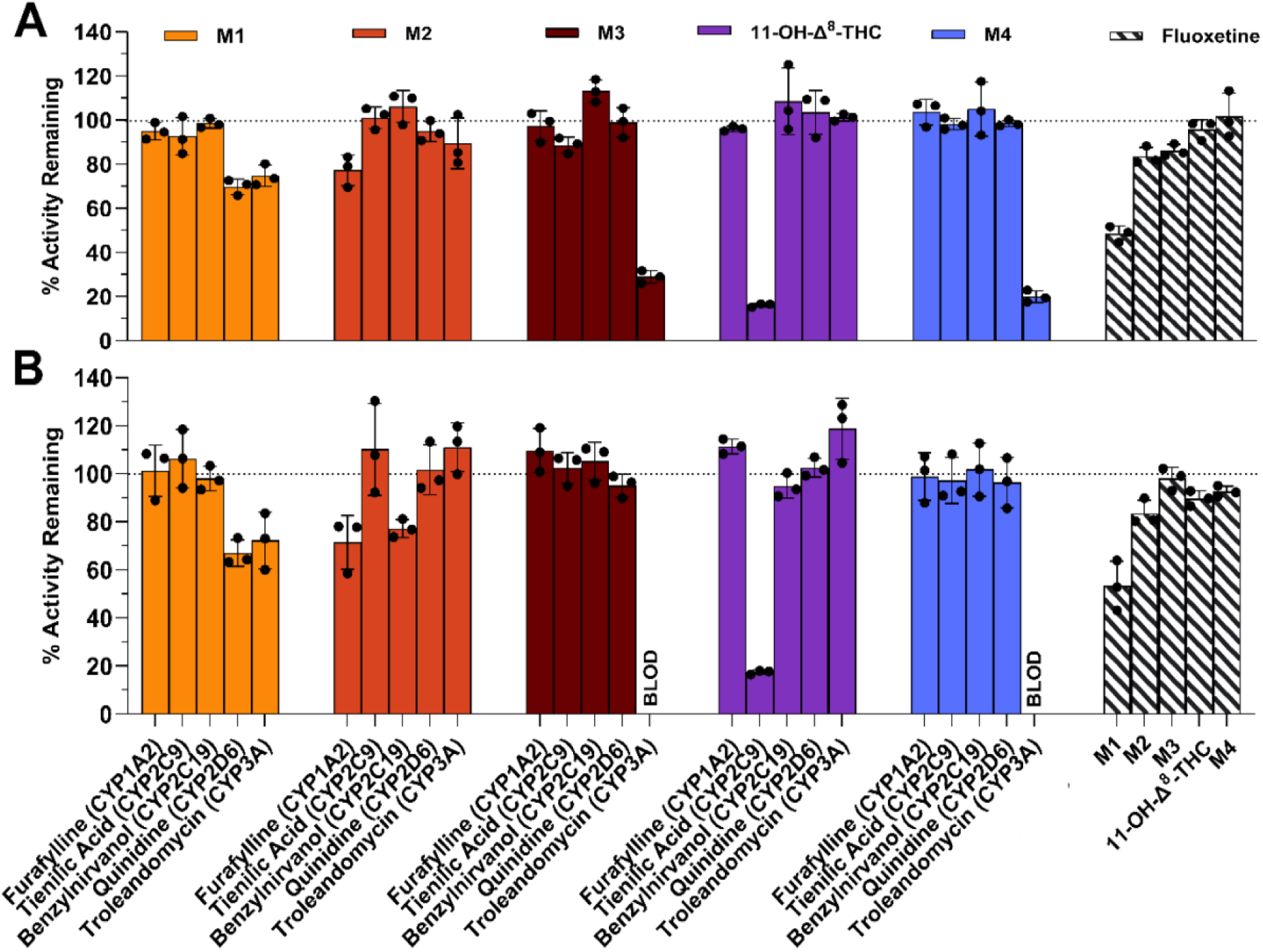
Inhibition of Δ^8^-THC metabolite formation in pooled human liver microsomes. The percent metabolite formation activity remaining relative to vehicle control in incubations of 2 µM (40 nM free concentration) (**A**) or 0.3 µM (6 nM free concentration) (**B**) Δ^8^-THC with selective CYP inhibitors (furafylline (CYP1A2), tienilic acid (CYP2C9), benzylnirvanol (CYP2C19), quinidine (CYP2D6), and troleandomycin (CYP3A) is shown for M1 (yellow), M2 (red), M3 (burgandy), 11-OH-Δ^8^-THC (purple), and M4 (blue). Slashed bars show the effect of fluoxetine on each of the metabolites. The dotted line at 100% indicates the uninhibited baseline. BLOD denotes metabolite formation below the limit of detection. The mean ± SD of three independent experiments is shown as the bars and symbols show the mean of duplicate incubations on each experiment.

Troleandomycin, a time dependent inhibitor of CYP3A, inhibited >70% of M3 and M4 formation in HLMs with product formation falling below the limit of detection at the lower substrate concentration (**Figure 6**). This suggests CYP3A is the main enzyme contributing to M3 and M4 formation. No other selective inhibitor produced meaningful reductions in M3 or M4 formation at either substrate concentration. For M1, quinidine (CYP2D6) and troleandomycin (CYP3A) reduced M1 formation by 25-34% (**Figure 6**). This suggests CYP2D6 and CYP3A both contribute to M1 formation in human liver. In contrast to quinidine, fluoxetine reduced M1 formation by 47-51% suggesting that in addition to CYP2D6, other CYPs inhibited by fluoxetine *in vitro*, such as CYP3A contribute to M1 formation in human liver. Inhibition of M1 formation by tienilic acid, benzylnirvanol and furafylline was negligible.

Several inhibitors reduced M2 formation. Furafylline inhibited M2 formation by 23-29% (**Figure 6**). Benzylnirvanol also reduced M2 formation by ∼26% at the lower unbound substrate concentration (6 nM), but no inhibition of M2 formation by benzylnirvanol was observed at the higher unbound substrate concentration (40 nM). The lack of inhibition at higher substrate concentrations is expected as benzylnirvanol is a competitive inhibitor, so its potency decreases at higher substrate concentrations. Substrate inhibition of CYP2C19 would further reduce the contribution of CYP2C19 at the higher concentration. Inhibition of M2 formation by fluoxetine was ∼20%, consistent with role of CYP2C19 in M2 formation. Overall, the inhibition experiments suggest M1 is formed by CYP2D6, CYP3A, and possibly CYP2C19, M2 is formed by CYP1A2 and CYP2C19, and M3 and M4 are formed by CYP3A. The formation of 11-OH-Δ^8^-THC is mainly by CYP2C9 with a minor contribution from CYP2C19.

To further evaluate the role of specific CYPs in metabolite formation, the formation of Δ^8^-THC metabolites was measured at 2 µM (free concentration 40 nM) substrate concentration in a panel of 22 individual HLM, and simple linear regression analysis was done between Δ^8^-THC metabolite formation rates and CYP-selective probe substrate metabolism (**Figure 7, Supplemental Table S4**). CYP1A2 and CYP2C9 activities significantly correlated with 11-OH-Δ^8^-THC formation, with CYP2C19 correlations showing a trend towards correlation. M1 formation correlated with CYP1A2, CYP2D6, and CYP3A activities. M2 formation correlated with CYP1A2 and CYP2C19 activities, and M3 and M4 formation each correlated with CYP3A activity. However, CYP2C9, CYP3A, and CYP1A2 activities, as well as CYP2C19 and CYP1A2 activities were co-linear across donors, confounding interpretation of these simple linear regression analyses. To account for this, multiple linear regression models incorporating all relevant CYP activities as covariates were implemented (**Table 2**).

**Figure 7:**
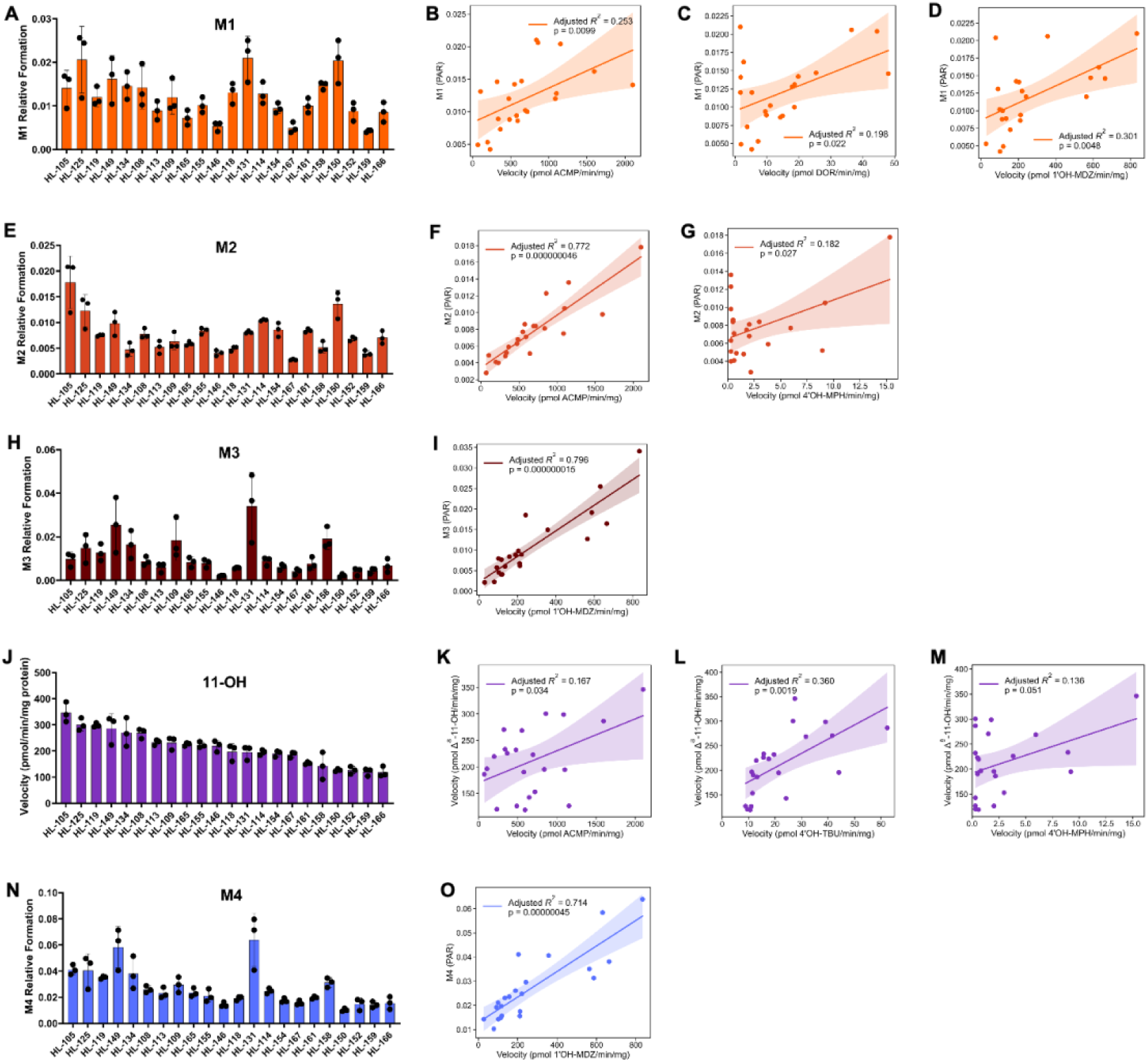
Δ^8^-THC metabolite formation in a panel of 22 human liver microsomes (HLM) and correlations with specific CYP activities. (**A**) M1 relative formation (peak area ratio, PAR) in a panel of HLM (rank-ordered by 11-OH-Δ^8^-THC formation velocity) and correlation between M1 formation and (**B**) CYP2D6 (dextrorphan, DOR) activity, (**C**) CYP1A2 (acetaminophen, ACMP) activity, and (**D**) CYP3A (1′-hydroxymidazolam, 1′OH-MDZ) activity. (**E**) M2 relative formation (PAR) in the same HLM panel and correlation between M2 formation and (**F**) CYP1A2 (ACMP) activity, (**G**) CYP2D6 (DOR) activity, and (**H**) CYP3A (1′OH-MDZ) activity. (**I**) M3 relative formation (PAR) in the same HLM panel and (**J**) correlation between M3 formation and CYP3A (1′OH-MDZ) activity. (**K**) 11-OH-Δ^8^-THC formation, expressed as absolute velocity (pmol min⁻¹ mg⁻¹ protein), in the same HLM panel and correlation between 11-OH-Δ^8^-THC formation velocity and (**L**) CYP2C9 (4′-hydroxytolbutamide, 4′OH-TBU) activity and (**M**) CYP2C19 (4′-hydroxymephenytoin, 4′OH-MPH) activity. (**N**) M4 relative formation (PAR) in the same HLM panel and (**O**) correlation between M4 formation and CYP3A (1′OH-MDZ) activity. For all bar graphs (**A, E, I, K, N**), bars represent the mean ± SD of duplicate incubations from three independent experimental days. For all correlation plots (**B–D, F–H, J, L, M, O**), solid lines represent the fitted simple linear regression and shaded bands the 95% confidence interval; each point represents a single HLM donor (n = 22). Adjusted R² and p-values are inset in each panel. Correlations with p < 0.05 were considered statistically significant. Analyses were performed in Python (v3.9.6). Incubations were performed at 5 µM Δ^8^-THC (free concentration 0.1 µM) as described in **Section 2.10**.

**Table 2:**
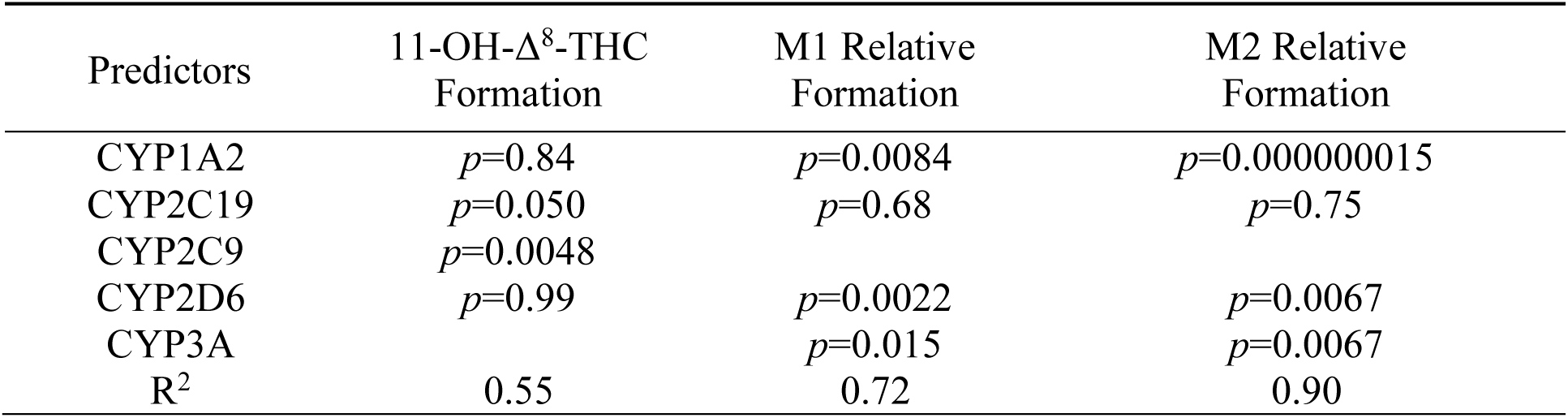
Multiple linear regression of Δ^8^-THC metabolite formation with CYP specific activities as covariates. p-values for each CYP isoform activity retained as a covariate in multiple linear regression models of 11-OH-Δ^8^-THC formation velocity or M1 and M2 relative formation are shown, with the R² of each full model listed in the bottom row (n = 22). Blank cells indicate predictors not included in the model for that metabolite. Predictors with p < 0.05 were considered significantly correlated with metabolite formation after adjusting for the other CYP activities.

In the multivariate models, CYP2C9 was the only significant predictor of 11-OH-Δ^8^-THC formation, with CYP2C19 approaching significance. CYP1A2, CYP2D6, and CYP3A activities were all significant predictors of M1 and M2 formation after adjusting for the other CYP activities, whereas CYP2C19 did not contribute to either model (**Table 2**). Variance inflation factors (VIF) for all predictors were below 5 in every model, indicating that residual multicollinearity was moderate and did not preclude interpretation of individual coefficients. The combined contribution of pairs of CYP activities to 11-OH-Δ^8^-THC, M1, and M2 formation was evaluated by three-dimensional multiple regression models relating two CYP-selective probe activities simultaneously to metabolite formation (**Figure 8**). For 11-OH-Δ^8^-THC, CYP2C19 and CYP2C9 activities were paired (**Figure 8A**). CYP2C9 was the stronger predictor of the two when modeled together. For M1, models pairing CYP1A2 with CYP2D6 (**Figure 8B**) and CYP3A with CYP2D6 (**Figure 8C**) explained a comparable proportion of the variability in formation velocity, with all three CYPs contributing significantly. For M2, models pairing CYP1A2 with CYP2D6 (**Figure 8D**) and CYP3A (**Figure 8E**) each provided a strong joint fit, consistent with the contributions of CYP1A2, CYP2D6, and CYP3A identified in the full multivariate model.

**Figure 8:**
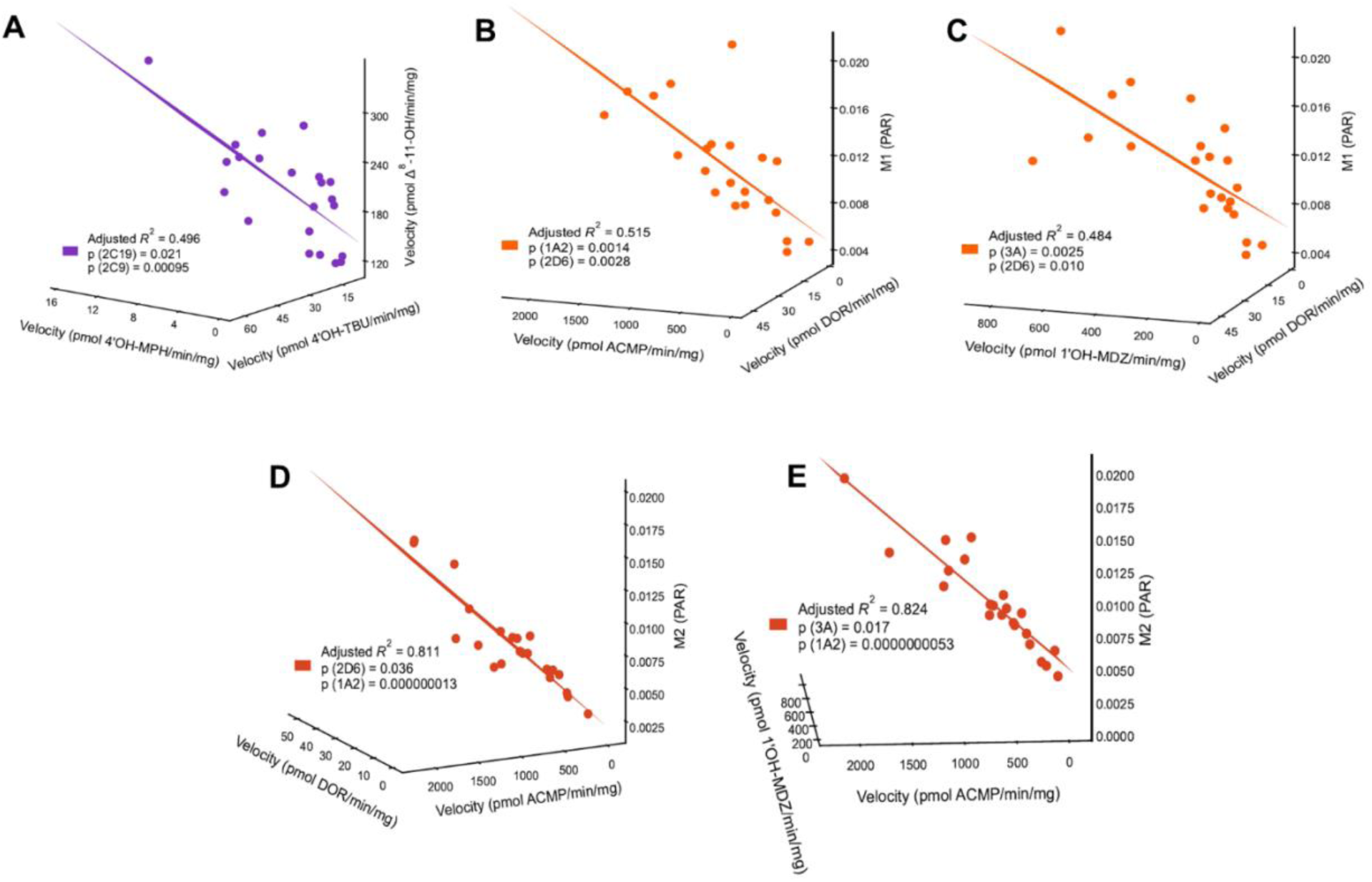
Multiple linear regressions relating paired CYP-selective probe activities to Δ^8^-THC metabolite formation in human liver microsomes (HLM). Three-dimensional multiple regression plots relate the formation velocities of two CYP-selective probe metabolites simultaneously to Δ^8^-THC metabolite formation across the 22 HLM donors. (A) 11-OH-Δ^8^-THC formation velocity regressed on CYP2C19 (4′-hydroxymephenytoin, 4′OH-MPH) and CYP2C9 (4′-hydroxytolbutamide, 4′OH-TBU) activity. (B) M1 formation velocity regressed on CYP1A2 (acetaminophen, ACMP) and CYP2D6 (dextrorphan, DOR) activity. (C) M1 formation velocity regressed on CYP3A (1′-hydroxymidazolam, 1′OH-MDZ) and CYP2D6 (DOR) activity. (D) M2 relative formation (peak area ratio, PAR) regressed on CYP1A2 (ACMP) and CYP2D6 (DOR) activity. (E) M2 relative formation (PAR) regressed on CYP3A (1′OH-MDZ) and CYP1A2 (ACMP) activity. Points represent individual HLM donors (n = 22), and the line shown is the tangent from the best-fit regression plane. Adjusted R² for the overall model and p-values for each individual predictor are inset in each panel. Predictors with p < 0.05 were considered statistically significant contributors to the model. Analyses were performed in Python (v3.9.6).

Nested linear regression models for 11-OH-Δ^8^-THC, M1, and M2 formation were further compared using partial F-tests (**Table 3**). M3 and M4 were not included in this analysis, as only CYP3A activity was consistently found to correlate with these metabolites. For 11-OH-Δ^8^-THC, addition of CYP2C19 significantly improved the CYP2C9 base model, whereas CYP1A2 and CYP2D6 provided no improvement. Addition of CYP1A2 and CYP3A significantly improved the M1 ∼ CYP2D6 model, consistent with the multiple regression results, whereas CYP2C19 did not. For M2 formation, adding CYP2D6 or CYP3A to a model containing CYP1A2 significantly improved fit, indicating that CYP2D6 and CYP3A activities explain some variance in M2 formation beyond that captured by CYP1A2 alone.

**Table 3:** Nested model comparison testing the contribution of additional specific CYP activity on Δ^8^-THC metabolite formation. Each base regression model (left column) was expanded by adding one CYP activity at a time as an additional predictor (middle column), and partial F test was performed. p-values (right column) indicate whether inclusion of the added predictor significantly improved model fit.

| Linear Regression | Added Predictor | *P-value* |
| --- | --- | --- |
| 11-OH-$\Delta^8$-THC ~ CYP2C9 | CYP1A2 | 0.34 |
|  | CYP2C19 | 0.021 |
|  | CYP2D6 | 0.64 |
| M1 ~ CYP2D6 | CYP1A2 | 0.0014 |
|  | CYP2C19 | 0.61 |
|  | CYP3A | 0.0025 |
| M2 ~ CYP1A2 | CYP2C19 | 0.54 |
|  | CYP2D6 | 0.036 |
|  | CYP3A | 0.017 |

### 3.4. Impact of genetic variability on Δ^8^-THC metabolism in human liver

CYP2C9, CYP2C19 and CYP2D6 were predicted to contribute to Δ^8^-THC metabolite formation in human liver. All three CYPs are subject to genetic polymorphisms that may impact the contribution of these CYPs to substrate metabolism. We hypothesized that genetic variability among the human liver donors underlies some of the interindividual differences in metabolite formation observed across the panel of HLMs. To test this hypothesis, we assessed the associations between CYP2C9, CYP2C19, and CYP2D6 genetic variability and Δ^8^-THC metabolite formation by Spearman rank correlation across the human liver panel (**Figure 9, Supplemental Table S5, Supplemental Figure S3**). CYP2C9 genotype based activity score was significantly associated with 11-OH-Δ^8^-THC formation (ρ = 0.46, p = 0.032) while neither CYP2C19 (ρ = −0.12, p = 0.61) nor CYP2D6 (ρ = −0.04, p = 0.85) genotype-based activity scores significantly associated with 11-OH-Δ^8^-THC formation (**Figure 9**). No significant correlation of CYP2C9 genotype-based activity score and M1–M4 formation was observed (ρ = −0.13 to 0.10) **(Supplemental Figure S3)**. A positive trend toward association between CYP2D6 activity score and M2 formation was observed (ρ = 0.37, p < 0.10), while no correlation was observed with M1 (ρ = 0.24). CYP2C19 activity score correlations were non-significant across all metabolites. (**Supplemental Figure S3, Supplemental Table S5**).

**Figure 9:**
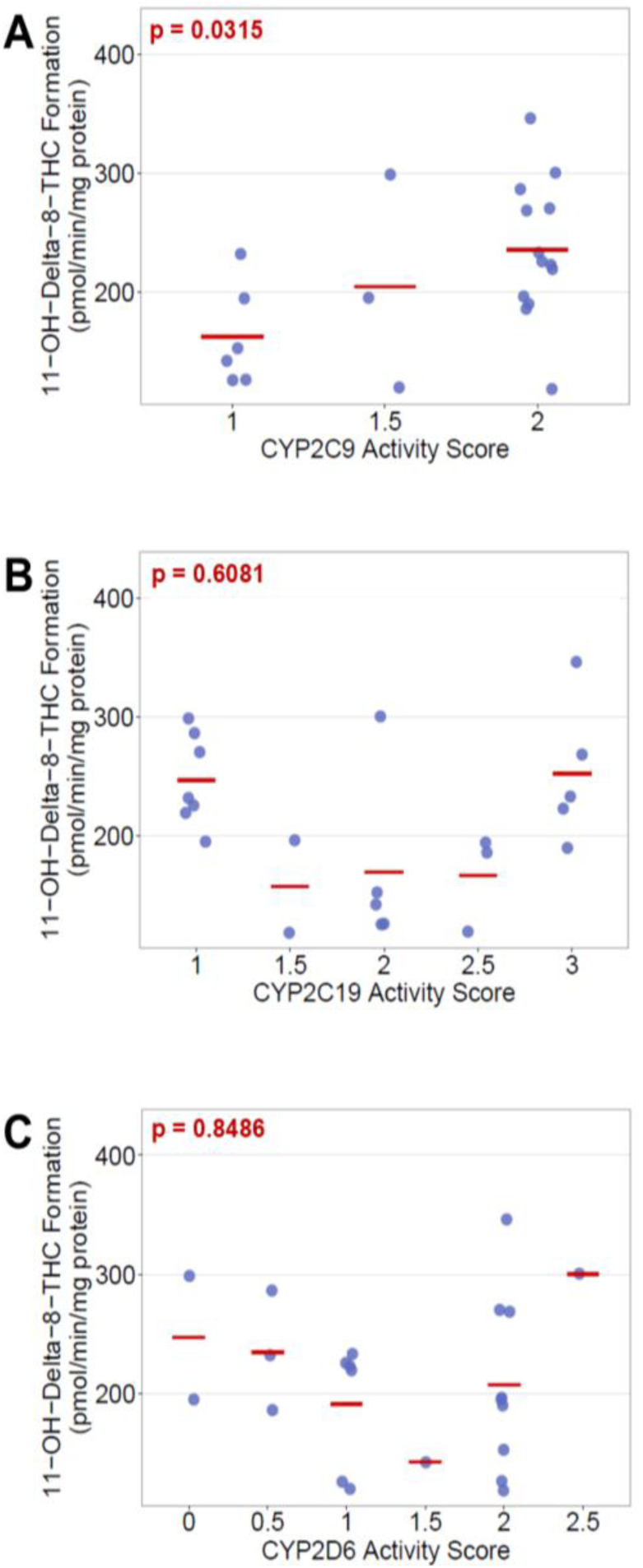
Associations of Δ^8^-THC metabolite formation with genotype-predicted CYP2C9, CYP2C19, and CYP2D6 activity scores. 11-OH-Δ^8^-THC formation (pmol min⁻¹ mg⁻¹ protein) stratified by genotype activity score in individual donors for CYP2C9 (**A**), CYP2C19 (**B**), and CYP2D6 (**C**) are shown. Each point represents an individual donor sample. Horizontal red lines indicate group means. P-values shown as inset are from Spearman rank correlation between activity score and 11-OH-Δ^8^-THC formation.

The impact of CYP2C9 genetic polymorphisms on 11-OH-Δ^8^-THC formation was further characterized using recombinant CYP2C9*2 and *3 Supersomes in comparison to the wild type CYP2C9*1 Supersomes. The 11-OH-Δ^8^-THC formation by CYP2C9*3 was significantly lower than by CYP2C9*1 and CYP2C9*2 (**Figure 10A**, **Table 4**). CYP2C9*1 showed the highest catalytic efficiency for 11-OH-Δ^8^-THC formation (CL_int,u_ = 162 µL min^-1^ pmol^-1^ CYP) followed by CYP2C9*2 (CL_int,u_ = 133 µL min^-1^ pmol^-1^ CYP). The efficiency of CYP2C9*3 was 94% lower (CL_int,u_ = 9.5 µL min^-1^ pmol^-1^ CYP). The 11-OH-Δ^8^-THC formation was scaled to microsomal formation clearance (**Figure 10B**). The predicted CYP2C9-mediated 11-OH-Δ^8^-THC formation clearance was 17,739 µL min^-1^ mg^-1^ protein in CYP2C9*1/*1 individuals, and the total 11-OH-Δ^8^-THC formation clearance was 14,706 µL min^-1^ mg^-1^ protein when CYP2C19 (2,377 µL min^-1^ mg^-1^ protein) was also accounted for. In CYP2C9*3/*3 individuals, the predicted CYP2C9-mediated 11-OH-Δ^8^-THC formation clearance was 670 µL min^-1^ mg^-1^ protein, and the total 11-OH-Δ^8^-THC formation clearance was 3,046 µL min^-1^ mg^-1^ protein, suggesting a 78% decrease in the overall 11-OH-Δ^8^-THC intrinsic formation clearance in CYP2C9 poor metabolizers.

**Figure 10:**
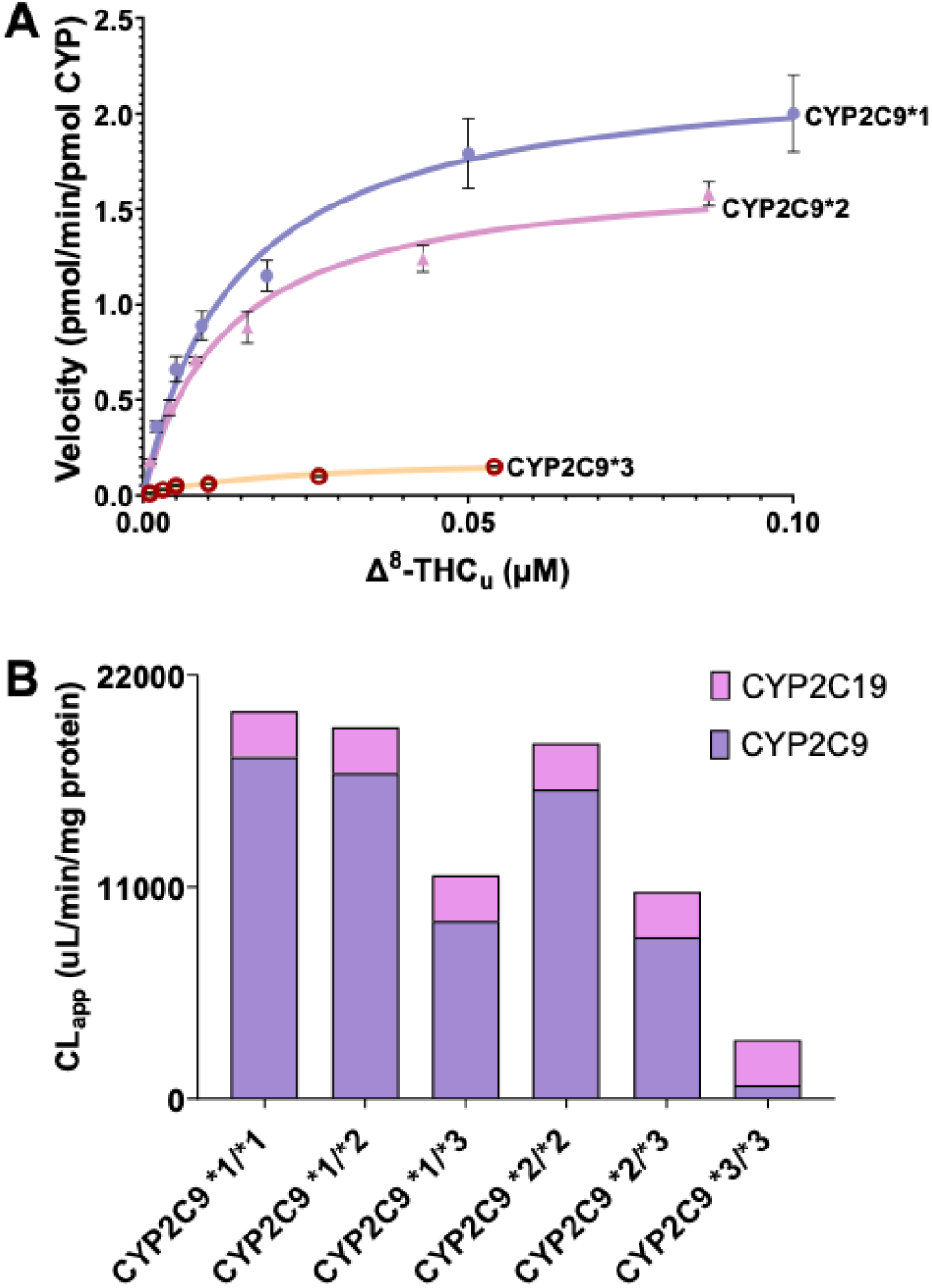
Effect of CYP2C9 genotype on 11-OH-Δ^8^-THC formation. **(A)** Michaelis–Menten kinetics of 11-OH-Δ^8^-THC formation by recombinant CYP2C9*1, *2, and *3 (velocity vs unbound Δ^8^-THC concentration; mean ± SD. (**B**) Predicted apparent intrinsic clearance (CL_app_, µL/min/mg microsomal protein) of 11-OH-Δ^8^-THC formation for the six diplotypes, scaled from recombinant activity. Bars are stacked by enzyme (CYP2C9, CYP2C19).

**Table 4:** Kinetic parameters for 11-OH-Δ^8^-THC formation by recombinant CYP2C9*1 (wild type), CYP2C9*2, and CYP2C9*3 Supersomes. . k_cat_, maximal formation rate; K_m,u_, unbound Michaelis constant; CL_int,u_, unbound intrinsic clearance (k_cat_/K_m,u_).

| **Parameter** | **CYP2C9\*1** | **CYP2C9\*2** | **CYP2C9\*3** |
| --- | --- | --- | --- |
| | $f_u = 0.16 \pm 0.08$ | $f_u = 0.14 \pm 0.08$ | $f_u = 0.08 \pm 0.05$ |
| $k_{\text{cat}}$ ($\text{min}^{-1}$) | $2.3 \pm 0.3$ | $1.7 \pm 0.06$ | $0.2 \pm 0.005$ |
| $K_{\text{m,u}}$ (nM) | $14 \pm 4.0$ | $13 \pm 1.2$ | $22 \pm 1$ |
| $\text{CL}_{\text{int,u}}$ ($\mu\text{L}/\text{min}/\text{pmol CYP}$) | $162 \pm 23$ | $133 \pm 12$ | $9.5 \pm 0.3$ |

## 4. Discussion

The goals of this study were to identify Δ^8^-THC metabolites formed by CYPs in human liver, define the enzymes catalyzing each pathway, and establish the impact of genetic polymorphisms in Δ^8^-THC metabolism in human liver. Five monohydroxylated metabolites of Δ^8^-THC were identified in HLM, and multiple CYPs contribute to their formation. Formation of the active 11-OH-Δ^8^-THC was dominated by CYP2C9 with a small contribution from CYP2C19. The C-ring hydroxylated metabolites M3 and M4 were assigned to CYP3A alone, and the pentyl side-chain hydroxylated metabolites M1 and M2 to CYP1A2, CYP2D6 and CYP3A4. These pathways expand on the previous report that only CYP2C9 and CYP2C19 catalyze the 11-hydroxylation of Δ^8^-THC.^31^ The findings on Δ^8^-THC metabolism have broad similarities with reported Δ⁹-THC metabolism,^16^ particularly C11 hydroxylation by CYP2C9 and C-ring hydroxylation by CYP3A. Some striking differences from Δ^9^-THC metabolism were also observed. For Δ^8^-THC, CYP2C19 efficiently catalyzed 11-OH-Δ^8^-THC formation while generating essentially no C-ring hydroxylated products, whereas with Δ^9^-THC the C-ring oxidation and C11 hydroxylation by CYP2C19 had similar efficiency.¹⁶ CYP1A2 and CYP2D6 in turn appeared more important for side-chain hydroxylation of Δ^8^-THC than reported for Δ^9^-THC.^16^

The *in vitro* findings in this study are consistent with 11-OH-Δ^8^-THC being the major metabolite of Δ^8^-THC and CYP2C9 being the dominant driver of Δ^8^-THC clearance in humans. Following Δ^8^-THC administration, 11-OH-Δ^8^-THC is the only primary Δ^8^-THC metabolite detected in plasma,^5,6^ corresponding to 11-OH-Δ^8^-THC being the major metabolite observed in HLM incubations. Further, the observed *in vitro* kinetics explain the differences in the *in vivo* pharmacokinetics of the active 11-OH-Δ^8^-THC and 11-OH-Δ^9^-THC metabolites. After equal oral doses of Δ^8^-THC and Δ^9^-THC, the 11-OH-THC-to-THC AUC ratio was lower after Δ^8^-THC dosing (0.5) than after Δ^9^-THC dosing (1.3).^5,6^ Interestingly, the CL_int,u_ for 11-OH-Δ^8^-THC formation in HLM (90 mL min^−1^ mg^−1^ protein) measured here is lower than that reported for 11-OH-Δ^9^-THC formation measured previously in the same pooled HLMs (372 mL min^−1^ mg^−1^ protein).^16^ Assuming plasma protein binding is similar for Δ^8^-THC and Δ^9^-THC, this difference in CL_int,u_ predicts 76% lower 11-OH-Δ^8^-THC formation *in vivo* than 11-OH-Δ^9^-THC and agrees well with the observed 61% lower 11-OH-THC-to-THC AUC ratio *in vivo*.

The lower CL_int,u_ of 11-OH-Δ^8^-THC formation in HLMs is likely mainly due to lower CL_int,u_ by CYP2C9. All the data shown here converge on CYP2C9 as the principal enzyme forming 11-OH-Δ^8^-THC in human liver. Scaling from recombinant enzymes predicted CYP2C9 f_m_ of 84%, tienilic acid inhibited 11-OH-Δ^8^-THC by ∼85% in HLM, and across the 22 donors, CYP2C9 activity and 11-OH-Δ^8^-THC formation correlated significantly. In addition, the K_m,u_ values for 11-OH-Δ^8^-THC formation in HLMs and with recombinant CYP2C9 were similar (3.6 and 8.8 nM, respectively).

However, the substrate inhibition kinetics observed in HLM and only with recombinant CYP2C19 point to a minor contribution of CYP2C19 to 11-OH-Δ^8^-THC formation, particularly at higher substrate concentrations. In addition to the substrate inhibition kinetics observed, the role of CYP2C19 is supported by the recombinant enzyme data which predicted a CYP2C19 contribution of 16% to 11-OH-Δ^8^-THC formation, and the correlation analysis in which addition of CYP2C19 activity improved the CYP2C9 base regression model.

The lower CL_int,u_ of 11-OH-Δ^8^-THC formation in comparison to 11-OH-Δ^9^-THC appears to be due to weaker affinity of Δ^8^-THC to CYP2C9. The K_m,u_ with recombinant CYP2C9 was 11-fold higher for Δ^8^-THC (8.8 ± 1.9 nM) than that reported for Δ^9^-THC (0.77 ± 0.16 nM).^16^ The K_m,u_ difference was smaller in HLM (3.6 nM for Δ^8^-THC versus 1.4 nM for Δ^9^-THC^16^) possibly due to additional enzyme contributions. The maximum catalytic rates of Δ^8^-THC and Δ^9^-THC C11 hydroxylation were within 2-fold of each other in HLM (325 ± 7 versus 521 ± 53 pmol min^−1^ mg^−1^ protein) and in CYP2C9 Supersomes (6.30 ± 0.19 versus 12 ± 1 min^−1^) with the Δ^8^-THC hydroxylation rate being consistently lower.

The relative importance of CYP2C9 in Δ^8^-THC clearance may be affected by substrate concentration and CYP2C9 genotype (**Figure 11**). The maximum unbound plasma concentration of Δ^8^-THC after consumption of 10 – 40 mg Δ^8^-THC via vaporization is 4–9 nM, assuming similar blood-to-plasma ratio and plasma f_u_ of Δ^8^-THC as reported for Δ^9^-THC.^5,32^ The K_m,u_ of Δ^8^-THC in HLM and CYP2C9 Supersomes (3.6 – 8.8 nM) is similar to the estimated maximum plasma unbound concentration of Δ^8^-THC. Hence some saturation of CYP2C9 *in vivo* is expected. Hepatic inlet concentrations after oral consumption of 10 – 40 mg Δ^8^-THC may also saturate CYP2C9, as the estimated maximum unbound hepatic inlet concentration after a 40 mg dose is 16 nM (assuming f_a_ = 0.45 and k_a_ = 0.7 h⁻¹ as for Δ^9^-THC,^33^ F_g_ = 1 and Q_h_ = 90 L/h) despite a maximum unbound systemic plasma concentration of only 0.4–2.6 nM.^6^ Typical doses of Δ^8^-THC in users are 10 – 40 mg,^34,35^ although heavy users may consume more. Total package content of oral products can reach 1,427 mg, with 75% of sampled products containing at least 87.5 mg.^34^ Hence CYP2C9 saturation and concentration dependent clearance of Δ^8^-THC is expected. Consistent with this saturable CYP2C9-mediated metabolism, oral Δ^8^-THC exposure (C_max_ and AUC) increased more than dose-proportionally across the 10–40 mg range, accompanied by a rise in the Δ^8^-THC:11-OH-Δ^8^-THC ratio at the highest dose.^6^ Upon saturation of CYP2C9, CYP2C19 becomes slightly more important in 11-OH-Δ^8^-THC formation (**Figure 11**). Higher contributions of other metabolic pathways via CYP3A4 and CYP1A2 are also expected due to the higher K_m,u_ values for these enzymes.

**Figure 11:**
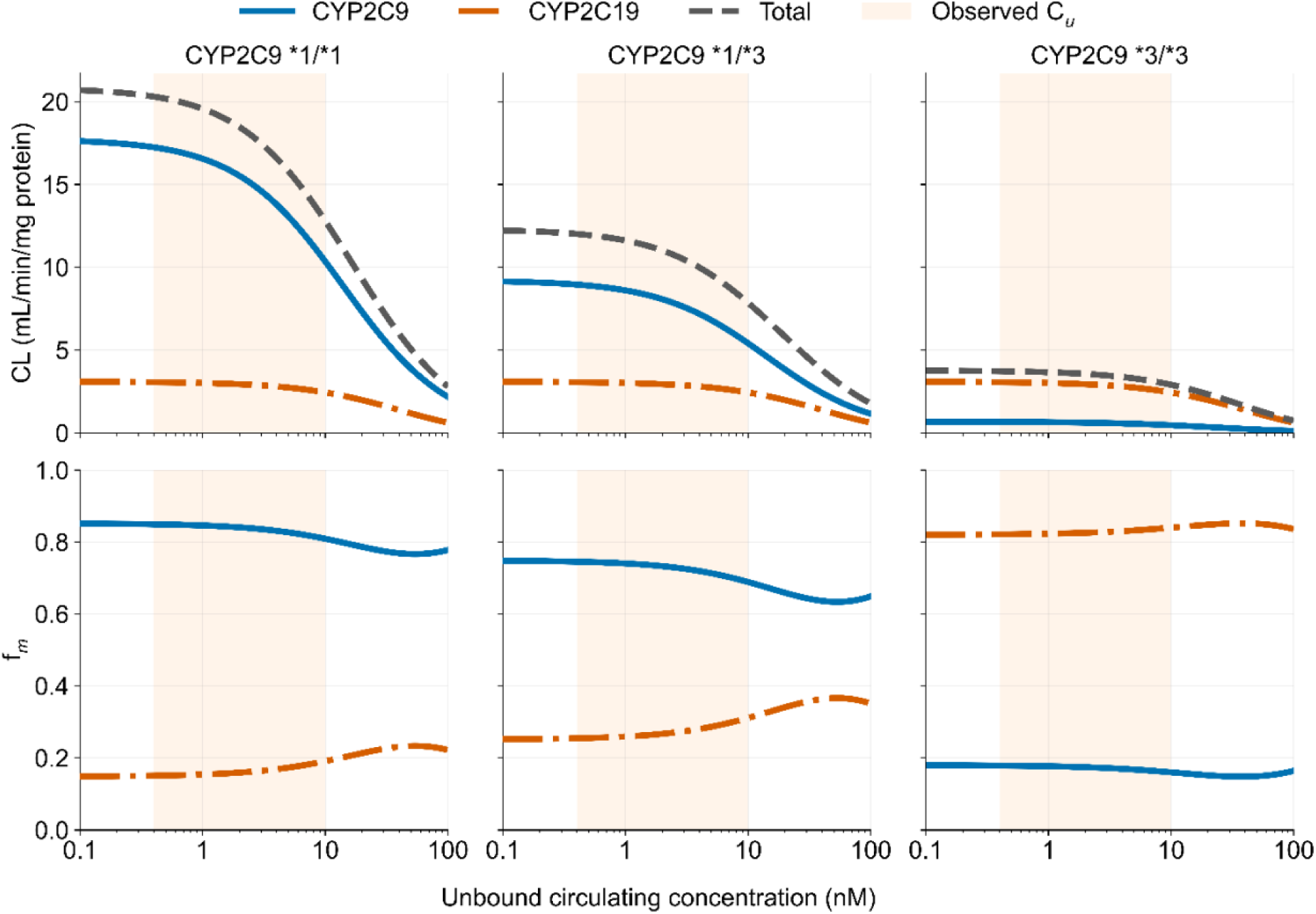
Predicted CYP2C9- and CYP2C19-mediated clearance and fractional contribution to 11-OH-Δ8-THC formation as a function of CYP2C9 genotype and unbound Δ^8^-THC concentration. Columns: CYP2C9 *1/*1 (left), *1/*3 (middle) and *3/*3 (right). Top row, enzyme-specific and total apparent clearance (CL, mL/min/mg microsomal protein); bottom row, fraction of total intrinsic clearance attributable to each enzyme (f_m_). x-axis, unbound circulating Δ^8^-THC concentration (nM, log scale, 0.1–100 nM). Solid blue line, CYP2C9; dash-dotted orange line, CYP2C19; dashed grey line, total (CYP2C9 + CYP2C19). Shaded band, range of observed unbound circulating concentrations (0.4–10 nM) reported following typical Δ^8^-THC exposures.

CYP2C9 saturation may also play a role in some of the *in vitro* experiments presented here. For example, the unbound concentration of 40 nM that was used in the high concentration inhibition experiments exceeds CYP2C9 K_m,u._ CYP2C9 f_m_ is expected to be slightly lower than that predicted from intrinsic clearance alone at the 40 nM unbound concentration (**Figure 11**), while CYP2C19 f_m_ is expected to be slightly higher. However, the impact of saturation on the overall Cl_int,u_ is predicted to be more striking than the impact on f_m_, likely due to the relatively low K_m,u_ for CYP2C19 as well (**Figure 11**). The inhibition experiments and correlation studies were consistent with the simulations based on the enzyme kinetics and expression levels (**Figure 11**). Tienilic acid inhibited 11-OH-Δ^8^-THC formation by ∼85% at both 40 and 6 nM unbound Δ^8^-THC concentrations, and CYP2C9 activity was a significant predictor of 11-OH-Δ^8^-THC formation velocity across the donor panel. Due to the high f_m,CYP2C9_, CYP2C9 genotype is expected to exert a considerable influence on Δ^8^-THC and the isoform contribution. In carriers of reduced-function CYP2C9 variants, both Δ^8^-THC clearance and CYP2C9 f_m_ are expected to decline, and in CYP2C9*3/*3 carriers the shift is sufficient that CYP2C19 is expected to become the predominant contributor to 11-OH-Δ^8^-THC formation (**Figure 11**).

In the in vitro assay results shown here, 11-OH-Δ^8^-THC formation CL_int,u_ by CYP2C9*3 was 6% of that by CYP2C9*1, while CYP2C9*2 activity was nearly identical to that of CYP2C9*1. The impact of CYP2C9*3 polymorphism is similar to a recent report of 11-OH-Δ^8^-THC formation by HEK cells expressing polymorphic CYP2C9*3, showing 98% reduced activity compared to CYP2C9*1. ^31^ However, the lack of effect of CYP2C9*2 observed here is in contrast to the prior report showing *2 and *3 having nearly identical activity towards Δ^8^-THC.^31^ The impact of CYP2C9*2 polymorphism observed here is also more modest than previously reported for Δ^9^-THC while the impact of CYP2C9*3 is similar.^36^ These differences between studies are likely due to differences in the expression systems used, and the normalization to reductase expression and correction to the measured unbound fraction done. The effect of CYP2C9*2 polymorphism on 11-OH-Δ^8^-THC formation observed here is similar to the lack of effect of CYP2C9*2 polymorphism on in vivo AUC of Δ^9^-THC.^14^ Similarly, Δ^9^-THC AUC is about 3-fold higher in CYP2C9*3/*3 carriers compared to CYP2C9*1/*1 carriers^14^ an effect size similar to what would be expected for Δ^8^-THC based on our data. Overall, as CYP2C9 is predicted to be the major Δ^8^-THC elimination pathway, Δ^8^-THC clearance and the f_m_ by CYP2C9 are expected to be lower in carriers of genetic CYP2C9 variants (**Figure 11**).

The interpretation of the data shown here is limited by the lack of authentic reference standards for M1–M4, and the positional assignments of the metabolites remain unconfirmed. Although linearity of the peak area response for all four new metabolites was verified by dilution series, the formation of these metabolites could be quantified only as a relative response. Absolute V_max_, k_cat,_ and CL_int_ values could not be determined for M1-M4. As a consequence, no mass balance could be established across the pathways, and the fraction of total Δ^8^-THC clearance attributable to each metabolite cannot be predicted. The study was confined to hepatic microsomal oxidation. Secondary metabolism of 11-OH-Δ^8^-THC to 11-COOH-Δ^8^-THC, further oxidation of M1–M4, and glucuronidation of all the compounds were not measured, nor was extrahepatic metabolism. We also did not test whether intracellular binding proteins such as FABP1 influence Δ^8^-THC metabolism. FABP1 alters both the rate of Δ^9^-THC oxidation and the relative CYP contributions in a metabolite and enzyme specific manner.^16^ The fractional contributions reported here may be impacted by FABP1 and therefore differ from those in intact hepatocytes.

In conclusion, this study shows that Δ^8^-THC is oxidized by human liver CYPs to at least five monohydroxylated metabolites, of which the active 11-OH-Δ^8^-THC is the principal product. CYP2C9 dominates its formation, with a minor contribution from CYP2C19 and the remaining pathways assigned to CYP3A, CYP1A2, and CYP2D6. 11-Hydroxylation is less efficient for Δ^8^-THC than for Δ^9^-THC. CYP2C9*3 polymorphism is expected to significantly reduce Δ^8^-THC clearance *in vivo,* similar to Δ^9^-THC. Overall, this study suggests that Δ^8^-THC disposition in humans is sensitive to CYP2C9 inhibition and CYP2C9 genotype and that pharmacological effects of Δ^8^-THC may in part be due to formation of active metabolites in humans.

## CRediT authorship contribution statement

K.T.S., Y.W.W: Conceptualization, methodology, validation, formal analysis, investigation, writing - original draft, writing - review and editing, visualization. Y.D., A.K.A., J.C., K.E.T: methodology, investigation, formal analysis, writing- review and editing N.I.: Conceptualization, investigation, writing - review and editing, supervision, funding acquisition.

## Acknowledgments

The authors thank Tim Creegan and David Stresser from Discovery Life Sciences for providing the polymorphic CYP2C9 Supersomes for this study.

## Nonstandard Abbreviations

1′-OH-MDZ: 1′-hydroxymidazolam
4′-OH-MPH: 4′-hydroxymephenytoin
4′-OH-TBU: 4′-hydroxytolbutamide
11-OH-Δ^8^-THC: 11-hydroxy-Δ^8^-tetrahydrocannabinol
11-OH-Δ^9^-THC: 11-hydroxy-Δ^9^-tetrahydrocannabinol
ACMP: acetaminophen
ANOVA: analysis of variance
AUC: area under the concentration– time curve
BLOD: below limit of detection
CI: confidence interval
CL_app_: apparent formation clearance
CL_int,u_: unbound intrinsic clearance
C_u_: unbound concentration
C_u,max_: maximal unbound concentration
CYP: cytochrome P450
Δ^8^-THC: Δ^8^-tetrahydrocannabinol
Δ^9^-THC: Δ^9^-tetrahydrocannabinol
Δ^9^-THC-COOH: 11-nor-9-carboxy-Δ^9^-tetrahydrocannabinol
DOR: dextrorphan
EIC: extracted ion chromatogram
FABP1: liver fatty acid binding protein
f_m_: fraction metabolized
f_u_: unbound fraction
h: Hill coefficient
HLM: human liver microsomes
IVIVE: *in vitro*–*in vivo* extrapolation
k_cat_: catalytic rate constant
K_i,u_: unbound substrate inhibition constant
K_m,u_: unbound Michaelis constant
LC–MS/MS: liquid chromatography–tandem mass spectrometry
LLOQ: lower limit of quantification
MRM: multiple reaction monitoring
m/z: mass-to-charge ratio
NADPH: reduced β-nicotinamide adenine dinucleotide phosphate
PAR: peak area ratio
POR: NADPH–cytochrome P450 oxidoreductase
S_50,u_: unbound half-maximal substrate concentration
SD: standard deviation
TAO: troleandomycin
VIF: variance inflation factor
V_max_: maximal formation velocity.

