## Supplemental materials for "Characterization of Cytochrome P450-Mediated Metabolism of Δ^8^-tetrahydrocannabinol *in vitro*"

**Title:** Cytochrome P450 Mediated Metabolism and Clearance Pathways of $\Delta^8$-THC

### 1. Methods

#### 1.1 Quantification of P450 oxidoreductase activity in supersomes of CYP allelic variants

NADPH–cytochrome P450 oxidoreductase (OR) activity in the CYP2C9 allelic variant supersomes (CYP2C9*1, *2 and *3 coexpressed with OR) and with purified human OR were measured spectrophotometrically using cytochrome c (equine heart, from Sigma, lot SLBQ2826V) as the electron acceptor. Assays were conducted in 100 mM potassium phosphate (KPi) buffer at pH 7.4 at room temperature. Cytochrome c working solution (360 µM) and fresh 5% (w/v) NADPH solution (50 mg/mL) were prepared in KPi buffer. Reaction mixtures were assembled in quartz cuvettes with a 1 cm pathlength to a final volume of 625 µL, containing 500 µL KPi buffer, 100 µL cytochrome c solution, and 25 µL CYP2C9. Samples were equilibrated in the spectrophotometer, and a baseline was recorded for 30 seconds. Reactions were initiated by addition of 25 µL NADPH solution, mixed directly in the cuvette. Final concentrations were 55.4 µM for cytochrome c, 77 nM CYP2C9, 1.9 mg/ml NADPH.

Spectral scans were performed on a Cary UV–Vis spectrophotometer (Agilent, Santa Clara, CA) equipped with Cary WinUV software. Kinetic monitoring was then performed in dual-wavelength mode at 550 nm (cytochrome c α-band) with reference at 541 nm. Instrument parameters included 0.2 absorbance full scale and a scan speed of 10 cm/min. Absorbance changes at 550–541 nm were recorded continuously for 3 min. A 1-min linear region of measured absorbance as a function of time, beginning immediately after NADPH addition, was used to determine the initial rate of cytochrome c reduction (ΔA/min) via linear regression in Cary WinUV kinetics software. Differences in these slopes were used to determine the relative OR content across the different CYP2C9 genotype lots. The OR activity units (AU) measure in nmol cytochrome c reduced per minute per microliter were 0.96 AU for CYP2C9*1, 1.77 AU CYP2C9 *2 and 2.37 AU CYP2C9*3. Based on these differences in measured OR activity, appropriate amount of purified OR was added to incubations with CYP2C9*1, *2 and *3 (OR coexpressed) supersomes to ensure each incubation had 5 activity units total of OR. Purified OR was incubated with the supersomes for 15 min at room temperature to allow reductase incorporation to the membrane. The incubations were then brought to a concentration of 100 nM CYP and used in the final incubation at 1.0 nM CYP2C9 per reaction.

**Supplemental Table S1: Intersystem Extrapolation Factors (ISEFs) used for hepatic scaling of $\Delta^8$-THC metabolite formation by recombinant CYP enzymes.** Literature reported ISEF values were used to scale apparent intrinsic clearance values ($CL_{int,app}$) measured in CYP Supersomes (BD Biosciences, Baculovirus expression system) to predict metabolite formation by each CYP in human liver microsomes (HLM). All reported ISEFs values were derived using Baculovirus-expressed Supersomes. Because no published ISEF for CYP3A5 is available, only CYP3A5 nonexpressors were considered in the analyses. $CL_{int}$, intrinsic clearance.

| CYP Enzyme | Probe Substrate (method obtained) | ISEF | Reference |
|---|---|---|---|
| **CYP1A2** | Phenacetin (substrate depletion) | 0.17 | Chen et al., 2011 |
| **CYP2C9** | Diclofenac ($CL_{int}$) | 0.54 | Chen et al., 2011 |
| **CYP2C19** | S-Mephenytoin (substrate depletion) | 0.25 | Chen et al., 2011 |
| **CYP2D6** | Dextromethorphan (substrate depletion) | 0.21 | Chen et al., 2011 |
| **CYP3A4** | Midazolam (substrate depletion) | 0.21 | Chen et al., 2011 |
| **CYP2J2** | 4-Hydroxyamiodarone ($CL_{int}$) | 0.0007 | Lee et al., 2011 |

**Supplemental Table S2: Population-Average Hepatic CYP Abundance Values Used for Predicting $\Delta^8$-THC Metabolism.** Hepatic CYP protein abundance values (pmol/mg microsomal protein) represent population-average estimates derived from published proteomic and immunoquantification studies of human liver microsomal preparations. CYP1A2 abundance represents the mean of values reported by Ohtsuki et al. (2012) and Kawakami et al. (2011); CYP3A4 and CYP3A5 population-average abundances were derived from Lin et al. (2002) but only CYP3A4 vaue was used due to the lack of ISEF value for CYP3A5.

| CYP Enzyme | Abundance (pmol/mg protein) | Reference |
|---|---|---|
| **CYP1A2** | 25.00 | Ohtsuki et al., 2012; Kawakami et al., 2011 |
| **CYP2C9** | 73.00 | Chen et al., 2011 |
| **CYP2C19** | 14.00 | Chen et al., 2011 |
| **CYP2D6** | 8.00 | Chen et al., 2011 |
| **CYP2J2** | 2.59 | Lee et al., 2011 |
| **CYP3A4** | 58.40 | Lin et al., 2002 |
| **CYP3A5** | 3.00 | Lin et al., 2002 |

#### 1.2 LC-MS/MS method for the quantification of dextrorphan and phenacetin

Acetaminophen and dextrorphan were quantified using a Sciex API 5500 QTRAP mass spectrometer (Framingham, MA) coupled to an Agilent 1290 Infinity II Ultra-High-Performance Liquid Chromatography system (Santa Clara, CA). Chromatographic separation was performed on an Agilent Zorbax Eclipse Plus C18 column (2.1 × 50 mm, 5 µm; Agilent Technologies, Santa Clara, CA) maintained at 35 °C, using mobile phases consisting of water with 0.1% formic acid (A) and acetonitrile with 0.1% formic acid (B). The mass spectrometer was operated in positive-ion electrospray ionization (ESI) mode with Turbo Spray, and analytes were detected by multiple reaction monitoring (MRM) with a dwell time of 100 msec per transition. Shared source parameters were as follows: ion spray voltage, 5500 V; source temperature, 600 °C; curtain gas, 35; collision gas, 7; and ion source gases 1 and 2, 50 each. Each analyte was quantified against a matched deuterated internal standard, and the two probe metabolites were analyzed using separate chromatographic gradients and compound-specific parameters, as described below.

For dextrorphan, 5 µL were injected. Mobile phase flow rate was 400 µL/min and the following gradient was used: 5% B held from 0 to 0.10 min, increased to 70% B at 3.50 min and to 95% B at 4.00 min, held at 95% B until 4.50 min, then returned to 5% B at 5.00 min and re-equilibrated until 6.50 min. Dextrorphan and its internal standard, dextrorphan-$d_3$, were monitored using the transitions *m/z* 258.0 → 157.0 and *m/z* 261.0 → 157.0, respectively, with identical compound parameters (declustering potential, 60 V; entrance potential, 10 V; collision energy, 51 V; collision cell exit potential, 20 V). For acetaminophen, 2 µL were injected. Mobile phase flow rate was 600 µL/min and the following gradient was used: 2% B held from 0 to 0.60 min, increased to 40% B at 2.00 min and to 95% B at 2.40 min, held at 95% B until 2.90 min, then returned to 2% B at 3.00 min and re-equilibrated until 4.50 min. Acetaminophen was quantified using the transition *m/z* 152.1 → 110.1 (collision energy, 30 V; collision cell exit potential, 12 V), with *m/z* 152.1 → 93.0 (collision energy, 30 V; collision cell exit potential, 10 V) monitored as a qualifier. Acetaminophen-$d_4$ was used as the internal standard (*m/z* 156.1 → 114.0; collision energy, 23 V; collision cell exit potential, 12 V). The declustering potential and entrance potential were 90 V and 8 V, respectively, for all acetaminophen transitions.

**Supplemental Table S3: Activity-score assignment for the different CYP2C9, CYP2C19, and CYP2D6 genotypes observed in the human liver bank.** For each enzyme, diplotypes (CYP2C9, CYP2C19) or allele function groups (CYP2D6) are listed with their corresponding activity score and predicted metabolizer phenotype. Activity scores were calculated as the sum of the two allele functional values for a given diplotype. CYP2C9 allele values: *1 = 1.0, *2 = 0.5, *3 = 0; CYP2C19 allele values: *1 = 1.0, *2 = 0, *17 = 1.5. CYP2D6 activity scores were calculated from the diplotype as reported previously (Shum et. al., 2022), with each allele assigned a functional value (normal = 1.0, decreased = 0.5, no function = 0) and the two values summed.

| CYP Enzyme | Genotype / Allele Function Group | Activity Score |
|---|---|---|
| **CYP2C9** | *1/*1 | 2.0 |
| | *1/*2 | 1.5 |
| | *1/*3 | 1.0 |
| **CYP2C19** | *17/*17 | 3.0 |
| | *1/*17 | 2.5 |
| | *1/*1 | 2.0 |
| | *2/*17 | 1.5 |
| | *1/*2 | 1.0 |
| **CYP2D6** | **Normal function**<br>*1, *2, *27, *33 | 1.0 per allele |
| | **Decreased function**<br>*14, *17, *29, *41, *59 | 0.5 per allele |
| | **No function**<br>*4, *5, *6, *56 | 0 per allele |

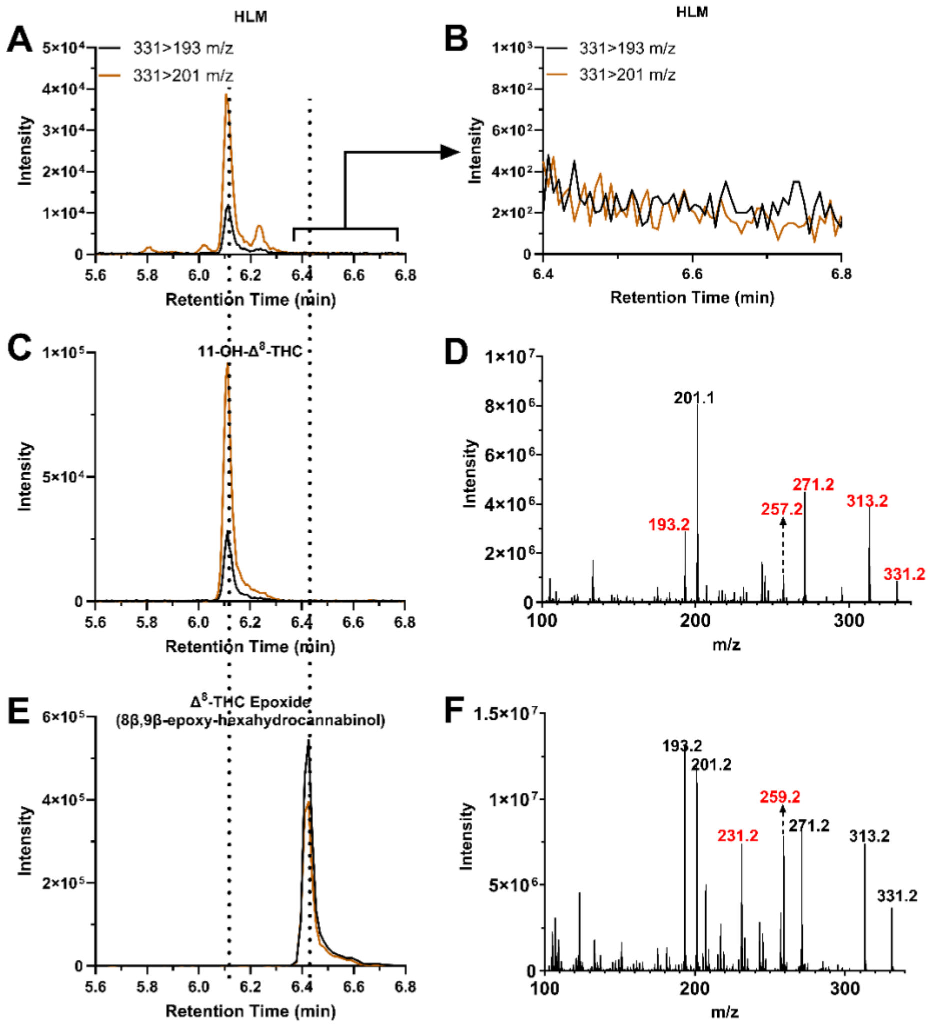


**Supplemental Figure S1: LC–MS/MS analysis of Δ8-THC metabolites generated by human liver microsomes (HLM).** (**A**) Extracted ion chromatograms (EICs) of the MRM transitions *m/z* 331→193 (black) and m/z 331→201 (orange), corresponding to oxygenated Δ8-THC metabolites ($[M+H]^+ = 331$), from an HLM incubation with Δ8-THC, showing a major peak at ~6.1 min and minor peaks indicating multiple oxidation products. (**B**) Expanded view of the boxed region in (**A**), displaying the same transitions from the HLM incubation over 6.4–6.8 min. (**C**) EICs of the same transitions for the 11-OH-Δ8-THC reference standard, which co-elutes with the major HLM peak at ~6.1 min. (**D**) MS/MS spectrum of 11-OH-Δ8-THC, dominated by *m/z* 201 with additional fragments at *m/z* 193, 257, 271, and 313, and the precursor at *m/z* 331. (**E**) EICs of the same transitions for the Δ8-THC epoxide (8β,9β-epoxy-hexahydrocannabinol) standard, which elutes later at ~6.4 min. (**F**) MS/MS spectrum of the Δ8-THC epoxide, dominated by *m/z* 193.2 and 201.2, with characteristic fragment ions at *m/z* 231.2, 259.2, 271.2, and 313.2, and the precursor at *m/z* 331.2.

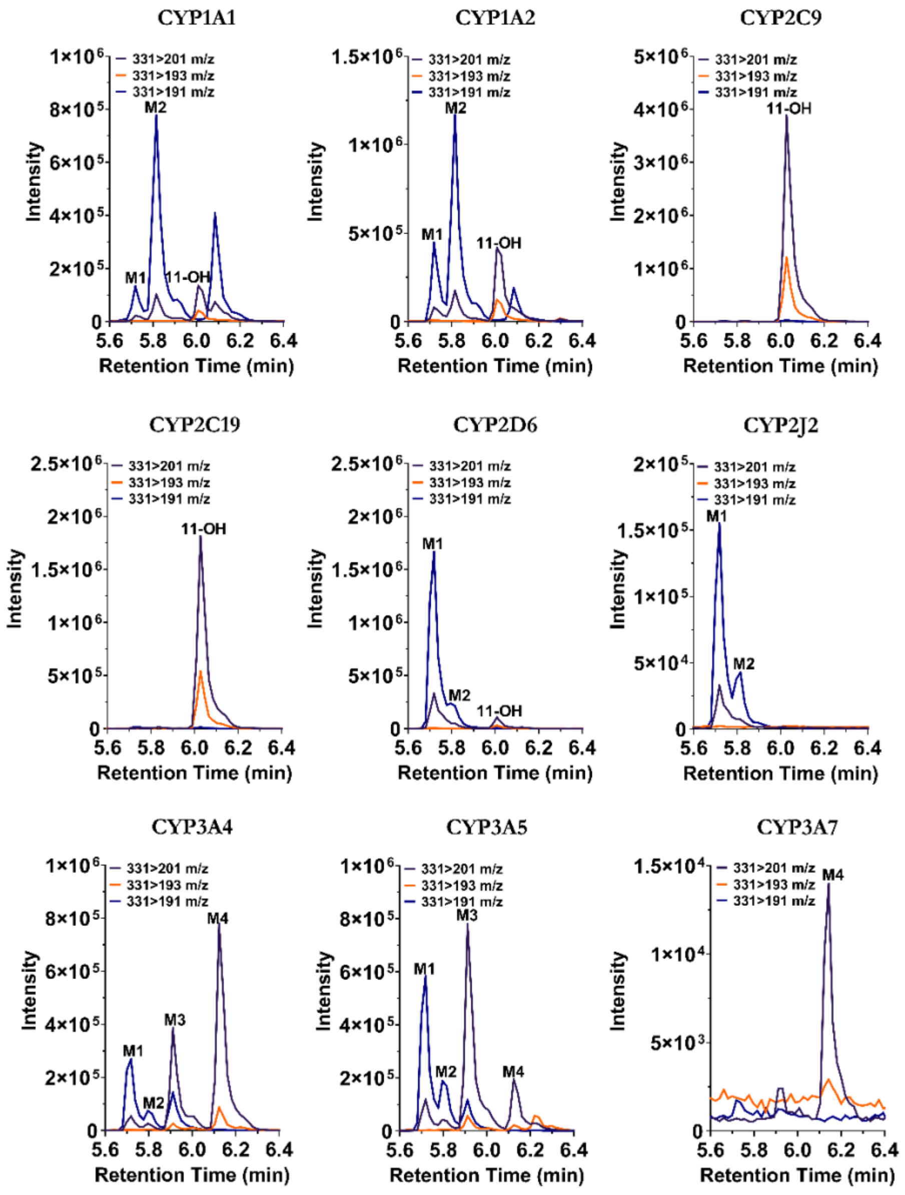


**Supplemental Figure S2:** Extracted ion chromatograms of the MRM transitions *m/z* 331→201, 331→193, and *m/z* 331→191 from incubations of Δ⁸-THC with individual recombinant CYP enzymes CYP1A1, CYP1A2, CYP2C9, CYP2C19, CYP2D6, CYP2J2, CYP3A4, CYP3A5, and CYP3A7. Labeled peaks indicate the metabolite(s) formed by each enzyme (M1–M4, 11-OH-Δ⁸-THC), illustrating the distinct metabolite formation profile of each CYP isoform.

**Supplemental Table S4: Statistics of the simple linear regression analyses of individual CYP specific activity against Δ⁸-THC metabolite formation.** p-values and $R^2$ for simple linear regressions between the formation velocity of each CYP-selective probe metabolite and 11-OH-Δ⁸-THC and M1–M4 formation velocity in human liver microsomes (n = 22). Correlations with $p < 0.05$ were considered statistically significant.

| | 11-OH-Δ⁸-THC Formation | M1 Relative Formation | M2 Relative Formation | M3 Relative Formation | M4 Relative Formation |
|---|---|---|---|---|---|
| CYP1A2 | $p$=0.034<br>$R^2$=0.21 | $p$=0.0099<br>$R^2$=0.29 | $p$=0.000000046<br>$R^2$=0.78 | | |
| CYP2C19 | $p$=0.051<br>$R^2$=0.18 | $p$=0.99<br>$R^2$=0.0000031 | $p$=0.027<br>$R^2$=0.22 | | |
| CYP2C9 | $p$=0.0019<br>$R^2$=0.39 | | | | |
| CYP2D6 | $p$=0.60<br>$R^2$=0.014 | $p$=0.022<br>$R^2$=0.24 | $p$=0.49<br>$R^2$=0.024 | | |
| CYP3A | | $p$=0.0048<br>$R^2$=0.33 | $p$=0.87<br>$R^2$=0.0014 | $p$=0.000000015<br>$R^2$=0.81 | $p$=0.00000045<br>$R^2$=0.73 |

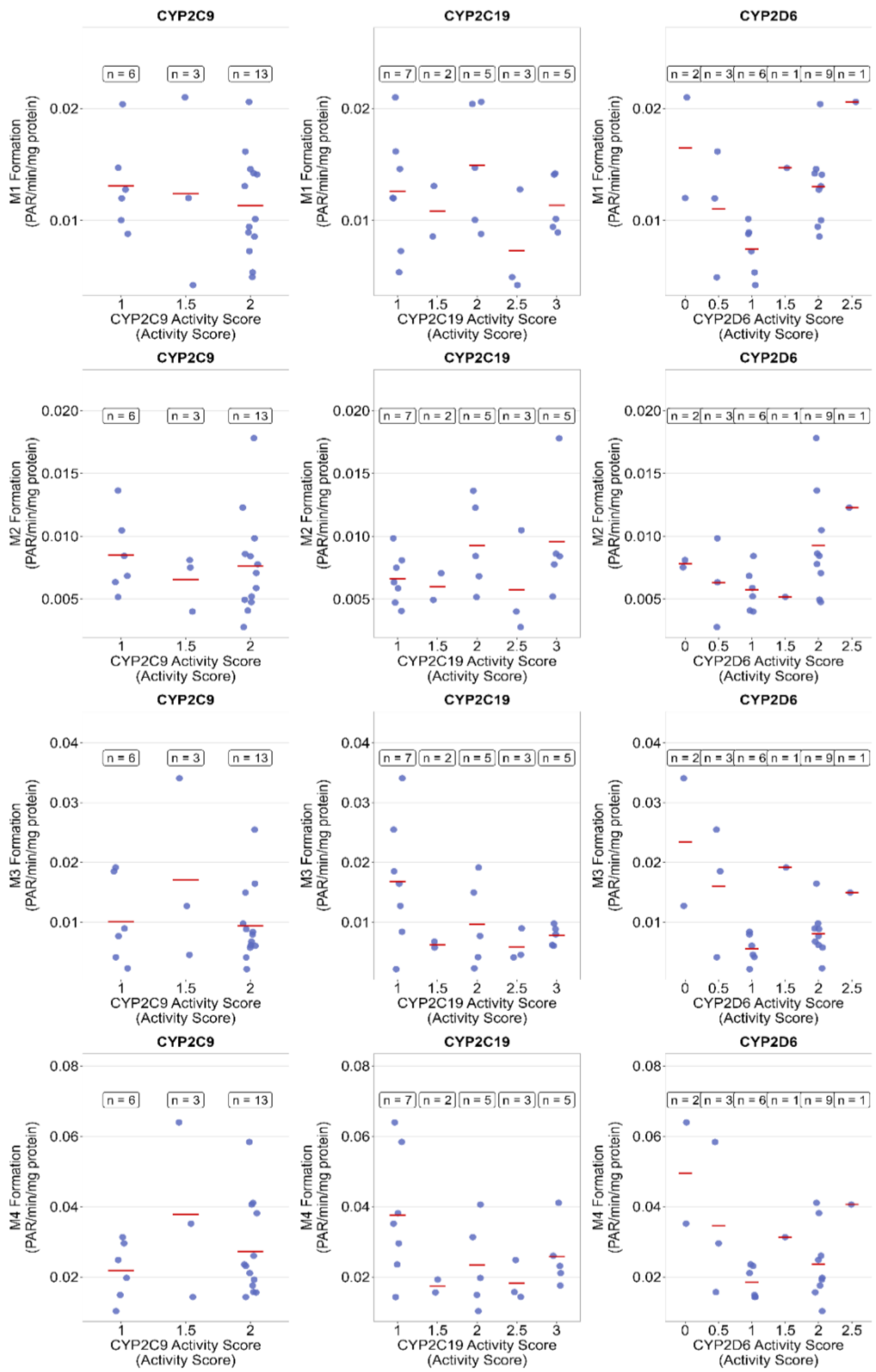


**Supplemental Figure S3: Association between CYP2C9, CYP2C19, and CYP2D6 genotype based activity scores and formation of Δ$^8$-THC metabolites M1–M4.** Formation of each metabolite (PAR/min/mg protein) is plotted against activity score for CYP2C9 (left column), CYP2C19 (middle column), and CYP2D6 (right column). Rows correspond to metabolites M1, M2, M3, and M4. Each point represents an individual donor sample, horizontal red lines indicate group means, and boxed values denote the number of donors per activity score group (n). Associations were evaluated by Spearman rank correlation and corresponding correlation coefficients and significance are summarized in **Supplemental Table S5**.

**SupplementalTable S5:** Spearman rank correlation coefficients (ρ) between CYP2C9, CYP2C19, and CYP2D6 genotype based activity scores and specific metabolic activity and the formation of $\Delta^8$-THC metabolites (11-OH, M1–M4).

| CYP enzyme | Metabolite | Spearman ρ | p-value |
|---|---|---|---|
| ***CYP2C9*** | 11-OH | 0.4594 | **0.0315** |
| | M1 | -0.1336 | 0.5533 |
| | M2 | -0.1259 | 0.5766 |
| | M3 | -0.0874 | 0.6990 |
| | M4 | 0.0996 | 0.6593 |
| ***CYP2C19*** | 11-OH | -0.1157 | 0.6081 |
| | M1 | -0.1535 | 0.4952 |
| | M2 | 0.2401 | 0.2817 |
| | M3 | -0.3233 | 0.1422 |
| | M4 | -0.2221 | 0.3205 |
| ***CYP2D6*** | 11-OH | -0.0432 | 0.8486 |
| | M1 | 0.2368 | 0.2886 |
| | M2 | 0.3694 | 0.0906 |
| | M3 | -0.1320 | 0.5581 |
| | M4 | -0.1196 | 0.5960 |
